# Identification and characterization of drug resistance mechanisms in cancer cells against Aurora kinase inhibitors CYC116 and ZM447439

**DOI:** 10.1101/2020.08.26.268128

**Authors:** Madhu Kollareddy, Daniella Zheleva, Petr Džubák, Josef Srovnal, Lenka Radová, Dalibor Doležal, Vladimíra Koudeláková, Pathik Subhashchandra Brahmkshatriya, Martin Lepšík, Pavel Hobza, Marián Hajdúch

## Abstract

CYC116 is a selective Aurora kinase inhibitor that has been tested in a Phase I study in patients with advanced solid tumors. Although CYC116 has shown desirable preclinical efficacy, the potential for emergence of resistance has not been explored. We established several CYC116 resistant clones from isogenic HCT116 p53+/+ and HCT116 p53−/− cell line pairs. We also generated resistant clones towards ZM447439 (quinazoline derivative), a model Aurora inhibitor. The selected clones were 10-80 fold resistant to CYC116 and cross-resistant to other synthetic Aurora inhibitors including AZD1152, VX-680, and MLN8054. Resistant clones displayed multidrug resistant phenotypes, tested by using 13 major cytostatics. All clones were highly resistant to etoposide followed by other drugs. Interestingly, all CYC116 clones but not ZM447439 became polyploid. ZM447439, but not CYC116 induced three novel mutations in Aurora B. Leu152Ser significantly affected ZM447439 binding, but not CYC116. Gene expression studies revealed differential expression of more than 200 genes. Some of these genes expression profiles were also observed in CYC116 resistant primary tumors. Bcl-xL (BCL2L1) was found to be overexpressed in CYC116 clones and its knockdown resensitized the p53+/+ resistant clone to CYC116. Finally Bcl-xL overexpressing p53+/+ CYC116 clones were highly sensitive to navitoclax (ABT-263) compared to parent cells. The data shed light on the genetic basis for resistance to Aurora kinase inhibitors which could be used to predict clinical response, to select patients who might benefit from therapy and to suggest suitable drug combinations for a particular patient population.

## Introduction

A major approach for effective cancer treatment in recent years is the development of targeted therapy. Focused research on biochemical pathways, involved in cancer genesis and progression, and evaluating differences between normal and transformed cells, allowed identifying new cancer targets. Few important examples of targets which are particularly involved in the transformation of a cell are cyclin dependent kinases (1), protein kinase B (Akt) (2), Bcl-2 (3), VEGFR-2 (4), B-RAF (5), BCR-ABL (6), and polo like kinases (7). Aurora kinases (AKs) (serine/threonine) have recently emerged as interesting drug targets. AKs A, B, and C are highly conserved and each has distinct and overlapping functions and sub-cellular locations. They are involved in the regulation of several multiple functions in mitosis including centrosome maturation, spindle assembly, chromosomal segregation, spindle mid-zone formation and cytokinesis. One of the hall marks of cancer cell is chromosomal instability. AKs regulate chromosomal stability at multiple levels. Several reports have been published that many cancers overexpress AKs, which leads to enhanced proliferation and chromosomal instability. AKs are widely considered as oncogenes (8).

Several Aurora kinase inhibitors (AKIs) are in various phases of anticancer clinical trials and some are in preclinical development (9). CYC116 ([4-(2-amino-4-methyl-thiazol-5-yl)pyrimidin-2-yl]-(4-morpholin-4-ylphenyl)-amine]) is a novel pan-Aurora kinase (Aurora A: 44 nM, B: 16 nM, & C: 95 nM IC50s) and VEGFR2 (IC50: 69 nM) inhibitor, which has been tested in a Phase I study. CYC116 induces mitotic failure and aneuploidy mainly due to inhibition of Aurora A and B kinases. It showed significant antiproliferative activity on various cancer cell lines and solid xenograft models (10).

CYC116 is a promising anticancer compound with significant specificity and potency, which could benefit patients with various cancers. Drug resistance is one of the main obstacles in successful cancer chemotherapy. Our study was focused on identification of potential cancer cell resistance mechanisms towards CYC116 alongside the first tool AKI, ZM447439 (11). Two isogenic colon cancer cell lines; HCT116 p53+/+ and HCT116 p53−/− were used as models to compare resistance mechanisms in the presence or absence of p53. All selected clones were highly resistant and also cross-resistant to other AKIs and to some approved cancer drugs. We found overexpression of antiapoptotic Bcl-xL (BCL2L1) in CYC116 resistant clones. Knockdown of Bcl-xL significantly restored the sensitivity to CYC116. Further, navitoclax (ABT-263), a small molecule Bcl-2 inhibitor (12) showed high activity selectively on Bcl-xL overexpressing p53+/+ resistant clones. These data suggest that the upregulation of Bcl-xL could limit the clinical response to CYC116. On the other hand, ZM447439 resistant clones acquired Aurora B mutations, which significantly affected drug binding.

## Materials and Methods

### Cell lines & Proliferation assay

HCT116 p53+/+ and HCT116 p53−/− cell lines were purchased from Horizon discovery. All cell lines were cultured in DMEM (Sigma) supplemented with 10% FBS, 100 μg/mL streptomycin, and 100 U/mL penicillin. MTT based proliferation assay was performed as described previously (13).

### AKIs and anticancer drugs

CYC116 was provided by Cyclacel Ltd. ZM447439 was purchased from Tocris, UK. AZD1152, VX680, and MLN8054 were purchased from Selleckchem. Anticancer drugs were purchased from Bristo-Myers Squibb (paclitaxel), Ebewe (doxorubicin, 5-fluorouracil, etoposide), Sigma (daunorubicin), Lachema (cisplatin, carboplatin, oxaloplatin), Lilly (gemcitabine), Unipharma (cladribine), Ovation (actinomycin-D), Glaxo (topotecan), Janssen-Cilag (bortezomib), and signalling inhibitors (ABT-263).

### Cell cycle analysis and pH3(Ser10) staining

The cell cycle analysis was carried out in three replicates as described previously (14). pH3 (Ser10) staining methodology can be found in the supplemental information.

### Computational modeling

Interactions of ZM447439 and CYC116 with the wild-type and several mutants of Aurora B kinase were studied using SQM/MM-based PM6-D3H4X method. Details of the computational methodologies are described in supplementary Information.

### Western blotting

Western blotting was performed as described previously (15). The membranes were probed with pH3Ser10 (06-570, Millipore), anti-Aurora A (N-20, Santa Cruz), anti-Aurora B (E-15, Santa Cruz), anti-p53 (BP53-12, Sigma), anti-Bcl-xL (clone 4A9, Origene), and β-tubulin (clone TU-06, Exbio) or β-actin (clone AC-74, Sigma) antibodies.

### DNA sequencing

Genomic DNA and RNA were isolated and cDNA was prepared as described previously (16). 25-50 ng of cDNA or gDNA was amplified by PCR using Phusion^®^ Hot Start High-Fidelity DNA Polymerase (Finnzymes) and specific primers. Approximately 50 ng of amplified cDNA (Aurora A, B, and C) and gDNA (Aurora B, C) were used in each sequencing reaction. Sequencing was performed on ABI Prism^®^ 3100-Avant Genetic Analyzer using Applied Biosystems chemistry. RidomTraceEdit (Ridom GmbH) and VectorNTI/ContigExpress (Invitrogen) softwares were used to align sequenograms and to check mutations. The primers used for sequencing can be found in supplemental Table S1.

### Gene expression and copy number studies and analysis

100 ng of RNA were used for preparation of biotinylated sense-strand DNA targets according to Affymetrix protocol. The fragmented and labeled sample was hybridized to Affymetrix Human Gene 1.0 ST Array. Expression profiles were examined from three independent biological replicates. All statistical analyses of expression arrays were carried out using either an assortment of R system software (http://www.R-project.org, version 2.11.0) packages including those of Bioconductor (version 2.7) by Gentleman et al. (17) or original R code. We used the affyQCReport Bioconductor R package to generate a QC report for all chips. Chips that did not pass this filter were not included. Raw feature data from the expression chips were normalized in batch using robust multi-array average (RMA) method by Irizarry et al. (18), implemented in R package affy. Based on the RMA log_2_ single-intensity expression data, we used Limma moderate T-tests (Bioconductor package limma (19) to identify differentially expressed genes. The p.adjust function from stats R package was used to estimate the false discovery rate using the Benjamini-Hochberg (BH) method (20). Expression data have been deposited in Array Express under accession number E-MEXP-3526. Parallel gene copy number analysis (array comparative hybridization) was evaluated in parent and resistant cells as described in supplementary information.

### Validation of differently expressed genes using qRT-PCR

The RNA was extracted using Trizol reagent (Ambion) followed by cDNA synthesis (Promega and Fermentas kits) and SYBR Green (Invitrogen) based qRT-PCR (Thermoscientific kit) (Rotor gene 6000 cycler). All gene primers were purchased from Generie Biotech. Thermal profiles were: 96° C for 15 m denaturation, then 95° C for 15 s amplification, and 66 or 64 or 62° C for 15 sec extension steps for 50 cycles. The specificity of gene primers and melting temperatures were optimized and the products were examined using Agilent DNA chips and analyzed using Agilent bioanalyser 2100. Target gene expression was normalized against to GAPDH housekeeping gene and 2^−ΔΔCt^ method was used to calculate the relative gene expression. Gene primers and thermal profiles are listed in supplementa1 Table S2.

### Validation of candidate genes using CYC116 *in vitro* sensitive and resistant primary human tumors

59 primary human tumor samples were used for analysis of *in vitro* response to CYC116 as described previously (21) and parallel tumor sample was snap frozen and stored at −80°C in tumor bank. 13 sensitive and 14 resistant tumors were selected for further validation study. The gene expression profiles were compared between CYC116 *in vitro* sensitive and resistant tumors. The Ct values were normalized against GAPDH. To calculate relative gene expression of resistant tumors in comparison to sensitive tumors unpaired t-test was used. These values were plotted in a chart to show relative gene expression differences between the sensitive and resistant tumors.

### siRNA mediated Bcl-xL knockdown

1.5 × 10^5^ cells were seeded in 6-well plates and incubated for 24 hours. Anti-Bcl-xL siRNAs (Origene) was diluted in jetPRIME (Polyplus transfection, France) buffer and transfection reagent, subsequently added to cells at a final concentration of 10 nM. After 24 hours of transfection, fresh media was added. Cellular lysates were prepared after 96 hours to follow the knockdown by western blotting. Among the three unique 27mer siRNA duplexes (named as A, B, and C), B and C types of siRNAs were very effective in depleting Bcl-xL. Combinations of B and C siRNAs at 5 nM each were therefore used for transfection followed by cell proliferation assay.

## Results

### Generation and selection of resistant clones

HCT116 p53+/+ and HCT116 p53−/− were exposed to cytotoxic concentrations (1 μM: above IC50) of CYC116 and ZM447439 in Petri dishes, which resulted in four groups of drug treated cell cultures: [**1]** p53+/+:CYC116, [**2]** p53−/−:CYC116, [**3]** p53+/+:ZM447439, and [**4**] p53−/−:ZM447439. During the course of selection, cells became polyploid, which is in agreement with Aurora B inhibition phenotype. Most of the cells died due to drug-induced effects. However, a small sub-population survived and formed colonies after 5 weeks. At least 10 colonies were isolated from each group and 3 clones were selected for further studies. Clones from each group were designated as: Group 1. [R1.1, R1.2, R1.3: CYC116 p53+/+], Group 2. [R2.1, R2.2, R2.3: CYC116 p53−/−], Group 3. [R3.1, R3.2, R3.3: ZM447439 p53+/+], and Group 4. [R4.1, R4.2, R4.3: ZM447439 p53−/−]. All clones were maintained continuously in 1 μM CYC116 or ZM447439 respectively.

### Clones displayed high resistance and cross-resistance to AKIs

p53+/+:CYC116 clones (R1.1, R1.2, R1.3) were 9, 82, and 62-folds less sensitive to CYC116 than the parent cells (Table 1). The differences in resistant factor values might be due to genetic heterogeneity within the clones in the course of selection process. Similarly p53−/−:CYC116 clones (R2.1, R2.2, R2.3) displayed high resistance to CYC116. Two clones from p53+/+:ZM447439 group exhibited very high resistance, >83-fold increase in IC50 values (R3.1, R3.2), where as the other clone (R3.3) is only 18-fold resistant. Degree of resistance was similar in p53−/−:ZM447439 group clones: 33, 38, and 39 folds, respectively. Overall, p53+/+ background displayed higher resistance to both AKIs.

**Table 1.** Resistance and cross-resistance profiles of parental versus CYC116 or ZM447439 resistant cells against various synthetic AKIs. All values in the table represent average IC50s in μM calculated from three independent experiments, each done in two technical replicates. The SD values for the above data were typically within 10-15% of the mean values. The values in parentheses are resistance factors.

| **Clone** | **CYC116** | **ZM447439** | **AZD1152** | **VX-680** | **MLN8054** |
| --- | --- | --- | --- | --- | --- |
| **HCT116 p53+/+** | 0.5 | 0.6 | 0.01 | 0.03 | 0.19 |
| **CYC116(p53+/+)** |  |  |  |  |  |
| R1.1 | **4.4 (9)** | 7 (12) | 17 (1700) | 1.9 (63) | 31 (163) |
| R1.2 | **41 (82)** | 12 (20) | 18 (1800) | 2.0 (67) | 15 (79) |
| R1.3 | **31 (62)** | 5 (8) | 11 (1100) | 1.0 (33) | 16 (84) |
| **HCT116 p53-/-** | 0.66 | 1 | >50 | 0.1 | 0.17 |
| **CYC116(p53-/-)** |  |  |  |  |  |
| R2.1 | **42 (64)** | 12 (12) | >50 | 4.0 (40) | 30 (176) |
| R2.2 | **27 (41)** | 7 (7) | >50 | 2.0 (20) | 3 (18) |
| R2.3 | **24 (36)** | 8.8 (8.8) | >50 | 2.4 (24) | 18 (106) |
| **ZM447439(p53+/+)** |  |  |  |  |  |
| R3.1 | 7 (14) | **>50 (>83)** | 36 (3600) | 2.6 (87) | 2.0 (10) |
| R3.2 | 1 (2) | **>50 (>83)** | 8 (800) | 0.7 (23) | 2.0 (10) |
| R3.3 | 1 (2) | **11 (18)** | 0.07 (7) | 0.09 (3) | 0.4 (2) |
| **ZM447439(p53-/-)** |  |  |  |  |  |
| R4.1 | 4.5 (7) | **33 (33)** | >50 | 0.8 (8) | 22 (129) |
| R4.2 | 3.0 (5) | **38 (38)** | >50 | 1.5 (15) | 18.6 (109) |
| R4.3 | 3.0 (5) | **39 (39)** | >50 | 3.0 (30) | 39 (229) |

Cross-resistance profiles were tested using AKIs that have been tested in clinical trials. p53+/+:CYC116 clones were highly cross-resistant to AZD1152 (1100-1800 fold) followed by MLN8054 (79-163 fold), and VX-680 (33-67 fold) (Table 1). Two clones (R2.1, R2.3) from p53−/−:CYC116 group were highly resistant to MLN8054 (176 & 106 folds) (Table 1). They were 20-24 fold more resistant to VX-680. AZD1152 was unable to reach the IC50 in p53−/− HCT116 cells - both parent and resistant clones even at the highest concentration tested (50 μM). p53+/+:ZM447439 clones presented variable resistance towards AZ1152 – from low to extremely high (7-3600 fold). They also displayed resistance to VX680 and MLN8054. p53−/−:ZM447439 clones were highly resistant to MLN8054 followed by VX680.

Interestingly, the p53−/−:ZM447439 cells were more resistant to MLN8054 than to the selecting agent. Overall, our data confirm wide cross-resistance among individual AKIs and suggest for shared mechanisms of drug resistance. All CYC116 and ZM447439 clones also acquired multidrug resistance phenotype as tested by using 13 approved anticancer drugs. Particularly all clones were highly resistant to etoposide. (supplemental Table S3). However, resistant clones were also more sensitive to several anticancer drugs compared to parent cell lines, particularly antimetabolites (5-fluorouracil and gemcitabine), DNA intercalators (daunorubicin and doxorubicin) and proteasome inhibitor bortezomib. Combination of these agents may have potential to prevent and/or overcome drug-induced resistance in concomitant or sequential administration.

### CYC116 clones, but not ZM447439 became polyploid irrespective of p53 status

Treatment of parent cells with either CYC116 or ZM447439 at a concentration of 1 μM for 48 h induced the mitotic failure and accumulation of G2/M and >G2/M cells in both p53+/+ and p53−/− cell lines (Fig. 1A-1J) and eventually apoptosis. Interestingly, all p53+/+:CYC116 clones have 4n (tetraploid) DNA content and p53−/−:CYC116 clones have slightly less than 4n DNA content (>3n-<4n) (Fig. 1C & 1D). On contrary to this, all ZM447439 clones were diploid (Fig. 1E & 1F). p53 levels in p53+/+ clones were equal or slightly up-regulated compared to controls (Fig. 1K).

**Figure 1.**
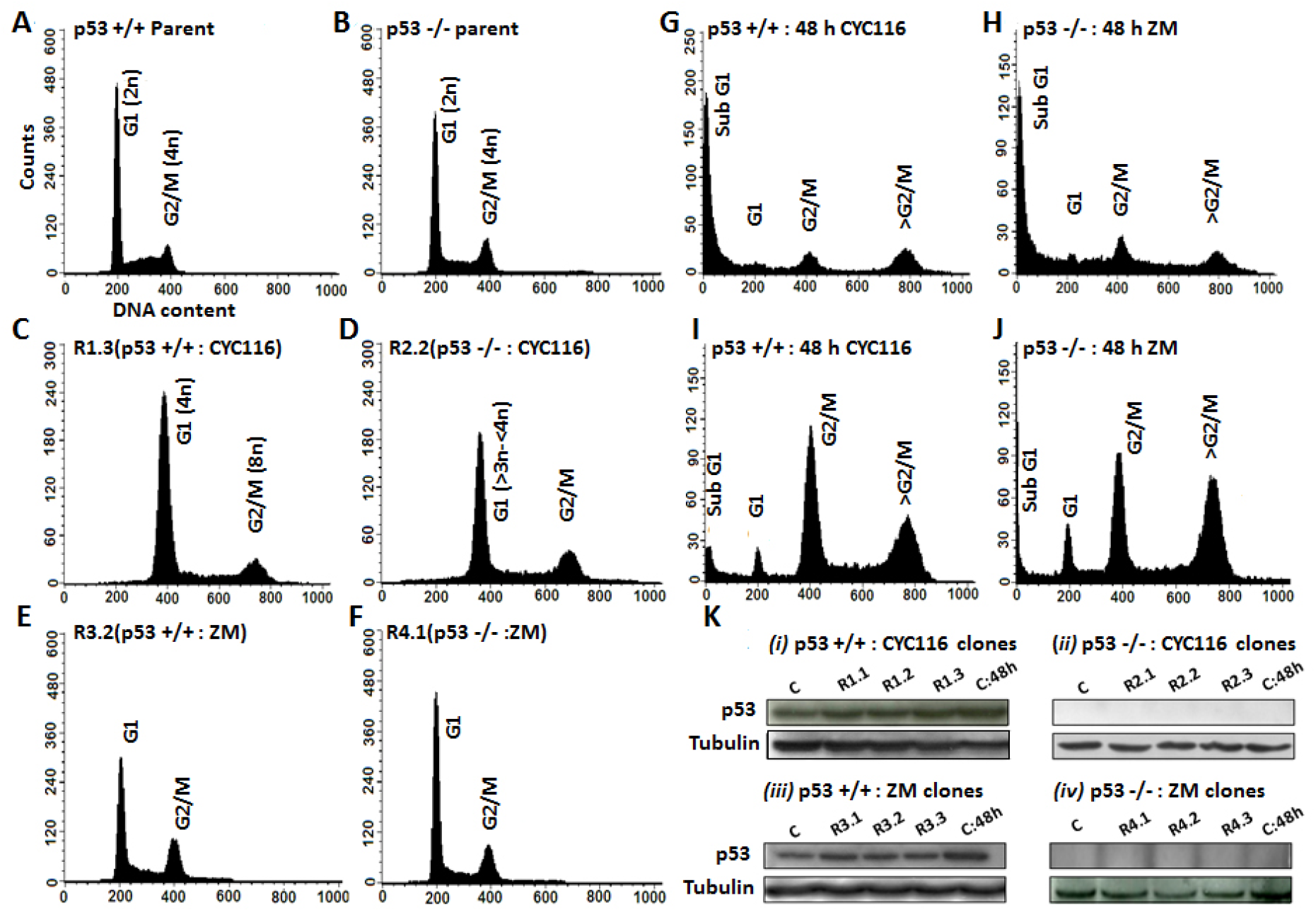
Cell cycle profiles of parent cell lines and resistant clones. A & B, Diploid parent HCT116 and HCT116 p53−/−. Cell cycle profiles for a representative clone form each group are shown. G, H, I, J, Cell cycle effects of CYC116 or ZM447439 on parent cell lines. C & D, Cell cycle profiles of CYC116 resistant clones. E & F, cell cycle profiles of ZM447439 clones. K, p53 induction levels in parent (C=DMSO: 48 h), resistant, and parent cell lines treated for 48 hours (C: 48 h).

### ZM447439 but not CYC116 induced mutations in Aurora B kinase and modeling of their impact on drug binding

DNA sequencing of AKs (A, B and C) revealed only three novel Aurora B mutations in ZM447439 clones (sequenograms in supplementary Fig. S1). One common mutation was detected in all p53+/+:ZM447439 clones i.e. I216L (mutant-1). Similarly one common mutation was detected in all p53−/−:ZM447439 clones, which is L152S (mutant-3). However R4.1 clone harbored L152S and one additional mutation, N76V (mutant-2). We carried out modeling of all three induced mutant proteins (mutant-1: I216L; mutant-2: N76V, L152S; mutant-3: L152S) to describe in structural and energy terms their effects on the interaction with ZM447439 and CYC116. Judging from the crystal structure of ZM447439 in complex with the wild-type Aurora B kinase (PDB code 2VRX) (22), the binding seems to be chiefly governed by dispersion (L83, L138, E125, L152, L154) with only one classical H-bond interaction (with A157) and one weaker C-H…O interaction with E155. The I216L mutation is far from the inhibitor (>7Å) and thus it does not directly influence the binding of the inhibitor (Supplemental Table S4). Further, it was seen that the side chain of I216 is buried in a pocket made up of I137, A187, L188, L184, L201, and L214. Our modeling study showed that I216L might lose some interactions (L184 & L214), which may clarify the resistance towards ZM447439. On the other hand, the terminal Cδ1 of L152 residue has three direct CH…π interactions with the inhibitor (23) (Fig. 2A). Thus its change to a smaller serine residue in both mutant-2 and mutant-3 is assumed to perturb these interactions. N76V is quite far from the binding site and hence it is beyond the scope of our study. However, it should be noted that this mutation disrupts the strong electrostatic interactions of Aspartate with T73, which might be one of the reasons for resistance.

**Figure 2.**
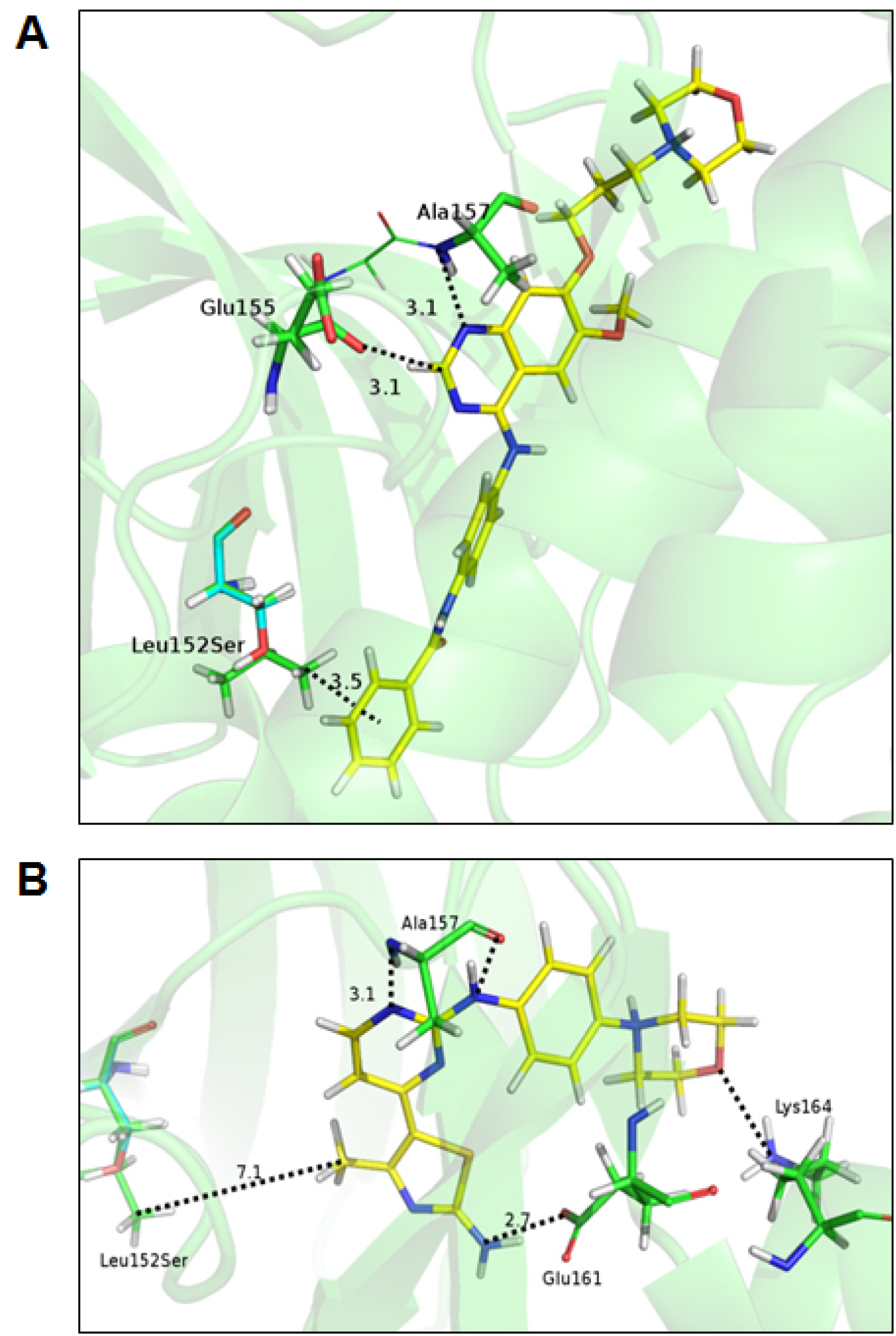
A close-up view of the binding interactions of ZM447439 and CYC116 in wild-type and in mutant-2 and 3 Aurora B kinases. A, The C-H…π interactions between L152 of Aurora B kinase and the terminal phenyl group ZM447439 are lost upon L152S mutation, which is not seen with CYC116. B, Specific hydrogen bond to the backbone of A157 and C-H…O H-bond to the backbone of E155 are shown along with two additional H-bond interactions (E161 and K164) in CYC116. Color coding: Ligand carbon in yellow, amino acids carbon in green, Ser152 carbon in cyan, nitrogen in blue, oxygen in red, hydrogen in white.

For CYC116, we performed docking to Aurora B kinase structure (PDB code 2VRX) and rescored the best docked pose. Interestingly, our docking results could identify identical interactions to those reported in previous studies which used a different docking program (10). CYC116 interacted differently with Aurora B kinase as compared to ZM447439 (Figure 2B). A prominent novel interaction was H-bond formed between the amino moiety of CYC116 and the side chain of E161. Similar to ZM447439, mutant-1 had a minor effect on the interactions as it is far (>7Å) from the active site. In contrast to ZM447439, L152 did not show any interactions with CYC116 as it was quite far from CYC116 (Figure 2B). Hence, we assume that CYC116 will not lose interactions against mutant-2 and mutant-3 proteins. The calculated binding energies between CYC116 and ZM447439 Aurora B mutants can be found in the supplementa1 Table S4.

Girdler et al. reported seven sets of Aurora B mutations induced by ZM447439 (22). Particularly G160E, Y156H, and G160V significantly affected ZM447439 binding. We performed modeling studies on these mutants using the CYC116 docked pose. We predict replacement of Gly with charged Glu might lead to some unknown quantum chemical phenomenon within the protein (e.g. electrostatic repulsion). This mutation also seemed to render cross-resistance to CYC116 significantly. For Y156H, in wild type (Y156), the phenyl ring of tyrosine showed π- π stacking interactions with pyrimidine ring of CYC116. Similarly, imidazole ring of histidine also interacted by stacking interactions and hence we assume that this mutation may not affect CYC116 binding. Like wild type G160, the mutant G160E conserved the CH… π stacking interactions with the aromatic ring of CYC116. Hence we believe that this mutation would not affect CYC116 binding. The binding energies can be found in the supplementa1 Table S4.

### Microarray based gene expression analysis

Whole human genome transcript array analysis was carried out to identify significant gene expression changes. The unsupervised clustering pattern suggest that majority of the gene expressions are common between the clones of each group. The clones were clustered (unsupervised clustering) with respect to p53 background and AKI used for selection of resistant clones (supplemental Fig. S2A). 50 genes were identified which mostly affect each of the first, second, and third component in PCA (principle component analysis) from all clones to generate the heat map (supplemental Fig. S2B). 885, 1085, 224, and 212 number of gene sets were differentially expressed (ANOVA p<0.001) in p53+/+:CYC116, p53−/−:CYC116, p53+/+:ZM447439, and p53−/−:ZM447439 groups, respectively. Compared to CYC116 clones, ZM447439 clones acquired small number of gene expression changes, which may suggest that ZM447439 clones acquired resistance predominantly by Aurora B mutations. The fold changes of all gene sets of each group and corresponding copy number changes were presented in the form of circular plots (Fig. 3). The top 100 genes from all four groups were listed (supplemental Tables S5A, S5B, S5C, S5D). Highly differentially expressed genes, common genes between the groups, and some based on biological relevance were further validated by qRT-PCR. Altogether 28 genes were selected from all groups (supplemental Table S6). Nearly 100% match and significant correlation was noticed between microarray data and qRT-PCR validation. (Supplemental Fig. S3).

**Figure 3.**
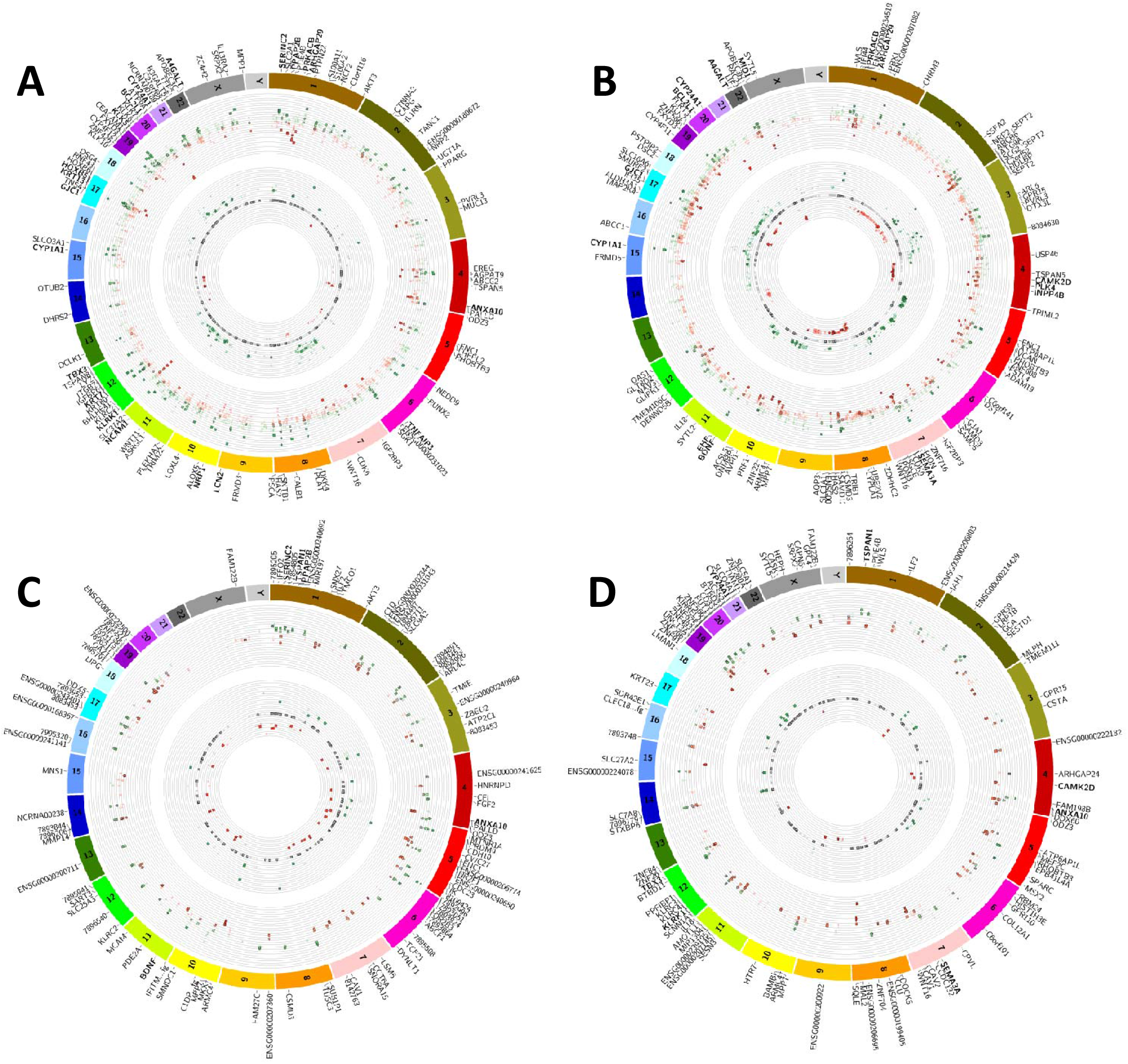
Circos visualization plots. The outer circular panel represents gene expression fold changes (logFC: fold change) of three clones from one group. Overexpression is color coded in green, whereas down-regulation in red. Grey color indicates no change in expression (−0.15 to 0.15). Inner circle represents corresponding gene copy number changes. Amplifications are color coded in green (>2 copies) and deletions in red (0-2). Grey color indicates no change in gene copy number. Each part of the circle coded in different colors represents chromosome number. Top 100 genes were highlighted in each group. The genes validated by qRT-PCR were marked as bold. Each clone was marked with a different symbol. A, HCT116: CYC116 (R1.1-□, R1.2-O, R1.3-Δ) B, HCT116 p53−/−:CYC116 (R2.1-□, R2.2-O, R2.3-Δ) C, HCT116:ZM447439 (R3.1-□, R3.2-O, R3.3-Δ) D, HCT116 p53−/−:ZM447439 (R4.1-□, R4.2-O, R4.3-Δ).

Since the number of differentially expressed genes is high, it is difficult to predict the affected pathways that influence AKIs induced resistance. We used GeneGo-system biology software to identify and prioritize most relevant pathways affected in resistant clones (Table 2). For HCT116 p53+/+:CYC116 clones the top scored common pathway map is apoptosis and survival-BAD phosphorylation (supplemental Fig. S4A). In HCT116 p53−/−:CYC116 clones the most relevant commonly affected pathway is retinol metabolism, where CYP1A1, CYP1B1, and CYP4F11 showed altered expression (supplemental Fig. S4B). The top common pathway that is affected in HCT116 p53−/−:ZM447439 clones is immune response-human NKG2D signaling (supplemental Fig. S4C). Finally the most relevant pathway affected in HCT116 p53−/−:ZM447439 is cell cycle regulation of mitosis (supplemental Fig. S4D). Some gene expressions which are altered in the above mentioned pathways were successfully validated by qRT-PCR (Bcl-xL, CYP1A1, PRKACB, KLRK1, and Cyclin H). Common or differentially affected pathways based on p53 background for CYC116 or ZM447439 resistant clones are shown in supplemental Table S7.

**Table 2.** List of commonly affected pathways in each group of resistant clones. ↓-down-regulation, ↑-up-regulation. Genes marked in bold indicate that have been validated successfully by RT-PCR.

| **Group** | **Commonly affected pathways and genes involved** |
| --- | --- |
| HCT116 p53+/+<br>CYC116 clones | Apoptosis and survival-BAD phosphorylation (**Bcl-xL**↑, **PKA-cat (PRKACB)** ↓, AKT3↑, GNG5 (G-protein beta/gamma)↓, PP2C (PDP1)↓ (supplemental Fig. S4A),<br>Development-PIP3 signaling (GNG5↓, AKT3↑, **Bcl-xL**↑, Cyclin D1↑),<br>Development-A3 receptor signaling (GNG5↓, AKT3↑, **PKA-cat** ↓, Cyclin D1↑),<br>Development-IGF-1 receptor signaling (IBP (IGFBP3 & IGFBP6)↑, **Bcl-xL**↑, AKT3↑, Cyclin D1↑,<br>PGE2 pathways in cancer (GNG5↓, GNAI1 (G-protein alpha-1)↓, AKT3↑, Axin2↑, **PKA-cat**↓, Cyclin D1↑ |
| HCT116 p53-/-<br>CYC116 clones | Retinol metabolism (**CYP1A1**↑, CYP1B1↑, DHA6↑) (supplemental Fig. S4B),<br>Cell-adhesion-Alpha-4 integrins in cell migration and adhesion ( FN1 (Fibronectin)↑, ITGB7↑, alpha-4/beta-7 integrin↑, **PKA-cat**↓,<br>Immune response-Antigen presentation by MHC class II (MHC class II↑, HLADRA↑) |
| HCT116 p53+/+<br>ZM447439 clones | Immune response-NK2D signaling (**KLRK1**↓, AKT↑, AP-1↓, c-Jun/c-Fos↓) (supplemental Fig.S4C),<br>Development ligand-dependent activation of ESR/AP1 pathway (RIP140↓, c-Jun/c-Fos↓),<br>Muscle contraction regulation of eNOS activity in endothelial cells (CAV1↓, AKT↑, c-Jun↓, ETV4 (PEA3)↑))<br>Reproduction-GnRH signaling (**PKA-cat** ↓, AP-1↓, c-Jun/c-Fos↓, AP-1↓) |
| HCT116 p53-/-<br>ZM447439 clones | Cell cycle-Initiation of mitosis (**Cyclin H**↓, APC↓, CAK complex↓, CDC25C↓, Weel↓) (supplemental Fig. S4D),<br>Cell cycle-Regulation of G1/S transition (Cak complex↓, Cyclin A↓)<br>Cholesterol and sphingolipids transport (CAV1↓, CAV2↓, ARH↑),<br>Histamine metabolism (ALDH2↑, MAOB↑, MAOA↑) |

### Comparison of drug resistant gene expression signatures in primary tumor biopsies sensitive/resistant to CYC116

Previously, several primary tumor samples (various solid and hematological cancers) were tested for sensitivity towards CYC116 under *in vitro* condition. 13 sensitive (average IC50: 4.42 μM) and 14 resistant samples (average IC50: 95 μM) were selected to validate gene expression signatures associated with response to AKIs. 23 most relevant genes were selected for qRT-PCR validation studies on human primary tumors *in vitro* sensitive/resistant to CYC116. Interestingly, majority of the cell line findings were also confirmed on primary human cells, suggesting validity of these genes as biomarkers of drug susceptibility or resistance (Fig. 4). Moreover, 5 genes expression (KRT7, PRKACB, EHF, ANXA10, CYP24A1) were statistically significant in this limited sample set. Our statistical analysis (receiver operating characteristic) showed that combination of these five genes serves as better predictive biomarkers of sensitivity towards CYC116 (in terms of both sensitivity and specificity) compared to any of these genes individually (supplemental Fig. S5)

**Figure 4.**
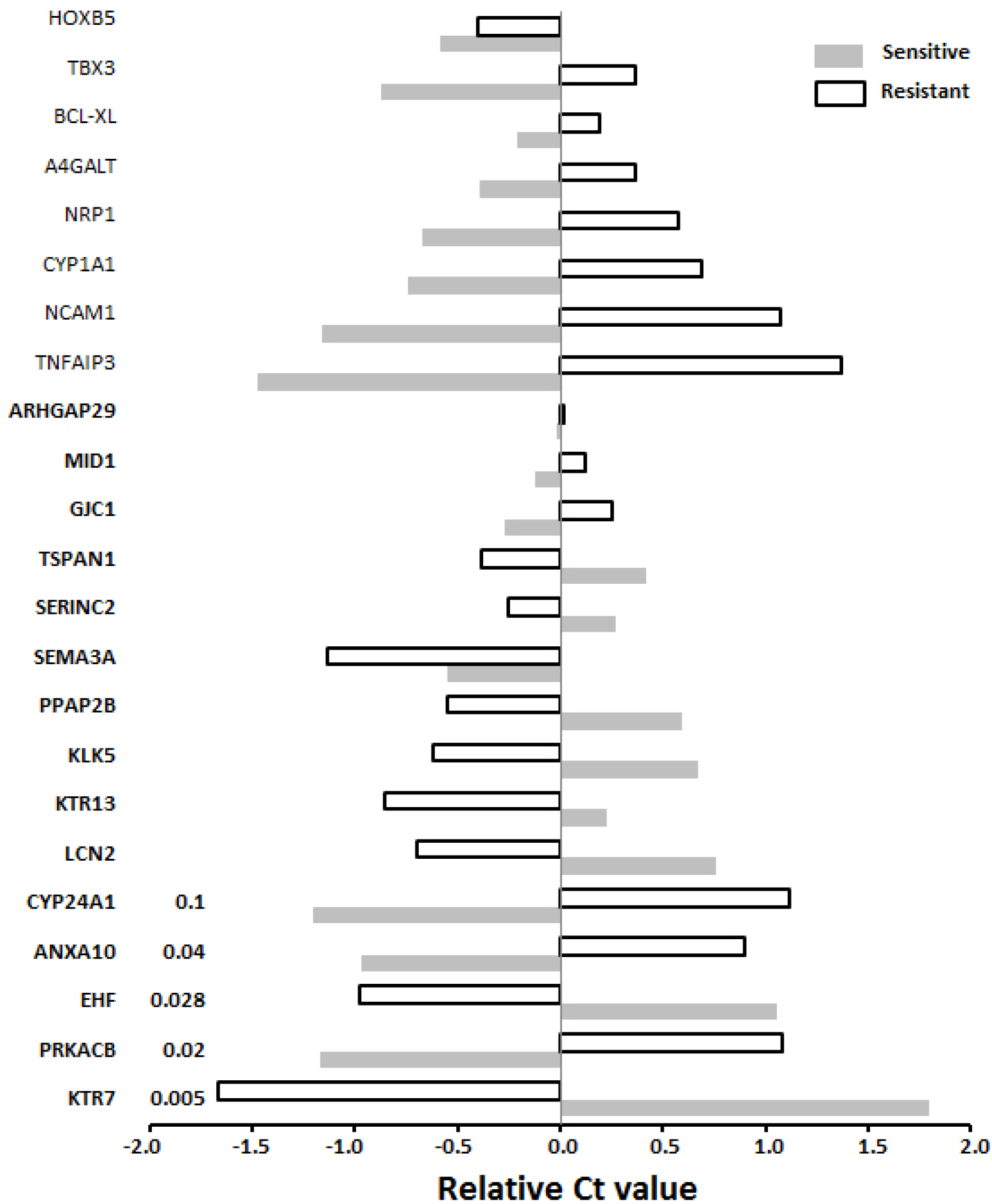
Validation of drug resistant gene expression signatures from cell lines in primary tumor biopsies sensitive/resistant to CYC116. The Ct value is reciprocal to the expression level. Gene expression trends that match to CYC116 resistant clones are marked in bold. Five genes expression trends shown from the bottom of the chart were statistically significant: KRT7, PRKACB, EHF, ANXA10, and CYP24A1. P-values for these five genes can be seen in the chart.

### Bcl-xL is overexpressed and its inhibition re-sensitized resistant clones towards CYC116

All CYC116 clones, but not ZM447439 showed upregulation of Bcl-xL at RNA and protein level. Expression of Bcl-xL in p53+/+ CYC116 resistant clones was much higher than p53−/− clones (Fig. 5A). Both B and C type of siRNAs (see Materials and Methods) were effective in depleting Bcl-xL(Fig. 5B). Cell proliferation/cytotoxicity assay was performed following Bcl-xL knockdown on two selected CYC116 resistant clones. Anti-Bcl-xL siRNA did not sensitize parent cells towards CYC116. Strikingly, depletion of Bcl-xL particularly in p53+/+ resistant clone significantly reversed the resistance. (Fig. 5C). We were able to reach the IC50 value at a level close to the sensitive parent cell line. Sensitization effect was much higher in p53+/+ than p53−/− background.

**Figure 5.**
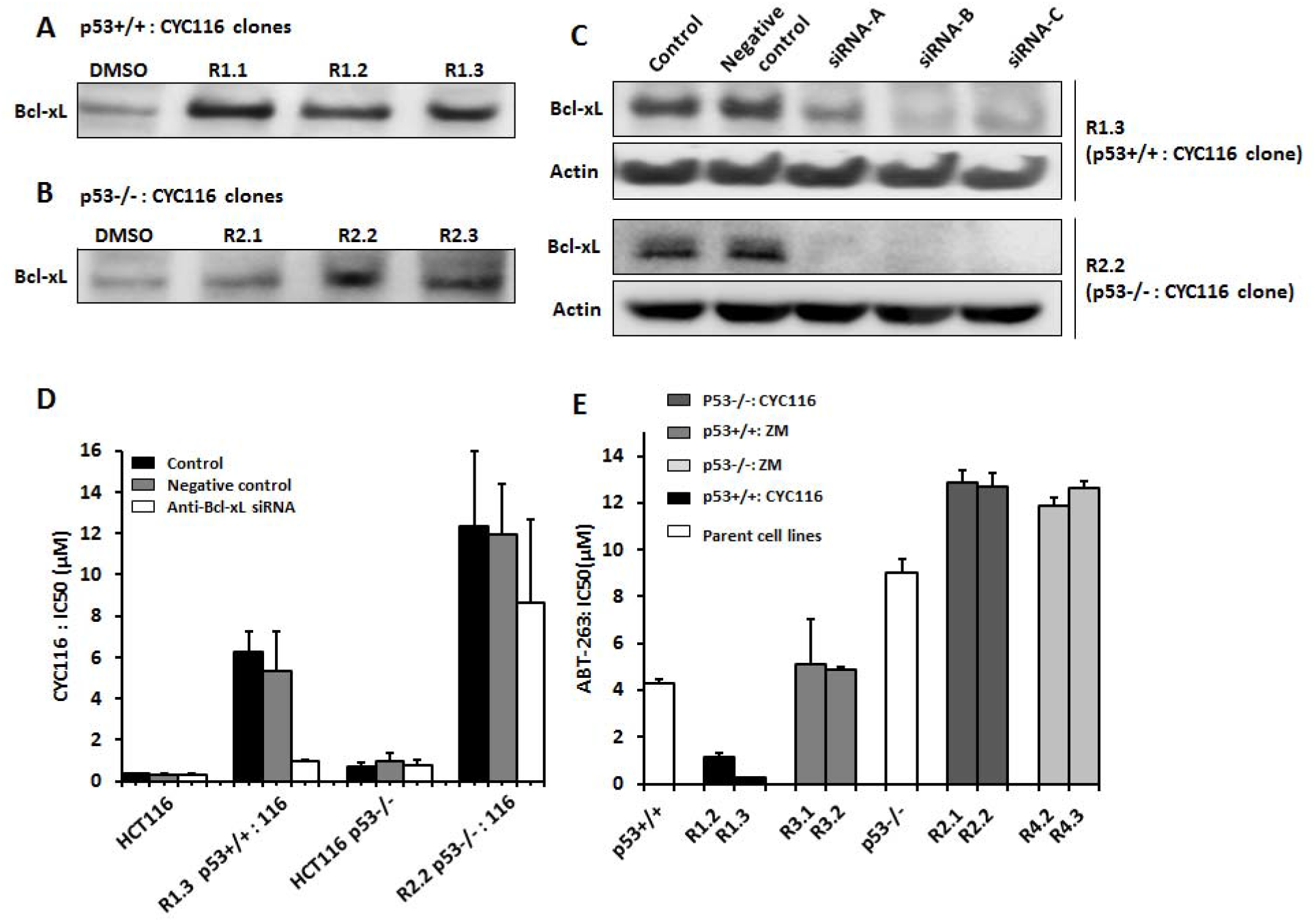
Inhibition of Bcl-xL expression restores response of CYC116 resistant cells to Aurora kinase inhibition. A, Bcl-xL expression in CYC116 resistant clones. Tubulin was used as a loading control as shown in Fig. 1 (same lysates were used for Bcl-xL expression also). B, Knockdown of Bcl-xL significantly by B and C types of anti-Bcl-xL siRNAs in R1.3 (p53+/+:CYC116) and R2.2 (p53−/−:CYC116) clones. C, IC50 values (n=3) of CYC116 after Bcl-xL knockdown in one p53+/+ & one p53−/− selected CYC116 resistant clones. D, IC50 values of ABT-263 on two selected resistant clones from each group (n=3).

### Bcl-xL expression correlates to the sensitivity of its pharmacological inhibition

Navitoclax (ABT-263) is a potent inhibitor of Bcl-2, Bcl-xL, and Bcl-w, which is currently in Phase II clinical studies in refractory cancers. Bcl-xL overexpressing CYC116 clones with p53 background were more sensitive (R1.2: 4 fold, R1.3: 17 fold) to ABT-263 than HCT116 parent cells (Fig. 5D). However, p53−/− CYC116 clones were resistant, suggesting a p53-dependent mode of action. All ZM447439 clones with no change in Bcl-xL expression were equally sensitive or slightly resistant to navitoclax. Cellular Bcl-xL levels significantly influenced the response to ABT-263 in p53+/+ CYC116 clones. In contrary, Bcl-xL expression in p53−/− cells (slightly up-regulated) did not modulate the response to ABT-263. This result was very much in line with siRNA mediated Bcl-xL knockdown, where Bcl-xL depletion sensitized p53+/+ CYC116 clone to a greater extent compared to p53−/− clone.

## Discussion

Understanding of genetic anomalies paved the way in identification of several targets and biomarkers specific to a cancer cell. This laid the rationale for so-called ‘personalized’ medicine and several targeted agents evolved and proved to be successful. Drug-induced resistance to these targeted agents is likely and has been evident in the clinic. Hence it is crucial to understand the genetic basis of resistance, which provides additional filter to stratify, select, and treat patients that would benefit from therapy.

Abnormal cell proliferation is one of the main hall marks of a cancer cell, which mainly depends on uninterrupted cell cycle progression. Several drugs were targeted to various essential nodes in the cell cycle machinery. AKs are one such entity, which are essential for cell cycle progression through mitosis. Aberrant expressions of AKs have been reported in several cancers (24–29), which formed the rationale for targeted therapy. CYC116 is a novel AKI with broad anticancer activity. We selected HCT116 to study CYC116-induced resistance, as they express little or no multidrug transporters (30), thereby reducing the chance of resistance due to drug pumps. Supporting this, flow cytometry based analysis of PgP and MRP1 expression revealed no induction in resistant clones (data not shown). Histone H3 is a direct downstream substrate of Aurora B, and inhibition of its Ser10 phosphorylation is a hallmark of Aurora B kinase inhibition (31). pH3Ser10 was not inhibited in resistant clones in the presence of AKIs as determined by flowcytometry and western blotting (supplemental Fig. S6). Western blotting revealed no significant changes in the expression levels of Aurora A and B in all resistant clones (supplemental Fig. S7). The CYC116 resistant and polyploid cell lines displayed altered cell cycle kinetics as evidenced by increase in doubling time to 1.2-2 folds. However all clones were actively cycling as evidenced by BrdU (DNA synthesis) and BrU (RNA synthesis) staining (data not shown). Further, the CYC116 resistant clones were highly proliferative, when xenografted into mice, suggesting that they did not lose tumorigenecity during the selection process (supplemental Fig. S8). Interestingly Bcl-xL overexpressing p53+/+ CYC116 clones showed enhanced tumor growth compared to p53 null clones. Chromosomal instability has been consistently reported to associate with MDR both in cell lines and patients. Castedo et al. established tetraploid HCT116 clones by treatment with other mitotic agents cytochalasin-D or nocodazole. They also selected tetraploid clones from diploid RKO cell lines by limiting dilutions (32). These clones were particularly resistant to DNA damaging agents including cisplatin, oxaloplatin, campothecin, and etoposide. However, inhibition or knockdown of p53 partially restored the cisplatin sensitivity. Our polyploid HCT116:CYC116 clones were also cross-resistant to some DNA damaging agents (supplemental Table S3), which is in line with their results. On the other hand, p53−/− CYC116 clones were either less resistant or sensitive to DNA damaging agents than p53+/+, which is again in agreement with their findings. Yuan et al. reported tetraploid clones in lenalidomide and bortezomib resistant multiple myeloma patient (33). Although this patient responded well initially, evolution of tetraploidy made resistant to lenalidomide and bortezomib with worsened prognosis. Lee et al. also reported worse prognosis in colorectal cancer patients with CIN+ (chromosomal instability) following 5-FU adjuvant therapy compared to CIN-tumors (34). Polyploidy per se provides survival advantage, because alterations in gene dosage can affect the target and drug stoichiometric ratios.

Our study also indicated that the CYC116 may be relatively ineffective in tumors that overexpress antiapoptotic Bcl-xL protein. The tumors which overexpress Bcl-xL may be also potentially insensitive to AZD1152, VX680, and MLN8054, as CYC116 clones are highly cross-resistant to these AKIs. Bcl-xL mediated drug resistance and relatively worse prognosis was reported consistently in the clinic (35). Here we report a novel resistance mechanism in the context of Aurora inhibition. Guo et al. found that SW620 and MiaPaca cell lines became resistant to AZD1152 (Aurora B inhibitor) by upregulation of PgP and BCRP, respectively (36). Seamon et al. showed upregulation of BCRP in JNJ-7706621 (Aurora A and B inhibitor) resistant HeLa cell line (37). Girdler et al. found several Aurora B mutations in ZM447439 (Aurora B specific) resistant HCT116 cell line, including H250Y, G160V, G160E, Y156H, and L308P (22). Further our collaborative study showed a range of protein candidates potentially involved in drug resistance against CYC116 (38).

Inhibition of Bcl-xL partially restored the sensitivity of resistant clones to CYC116; suggesting involvement of additional mechanisms. Indeed our genego analysis showed few relevant pathways affected in resistant clones including anti-apoptotic survival and drug metabolism pathways. The pathway analysis predicted the regulation of Bcl-xL by AKT via BAD phosphorylation in HCT116 p53+/+: CYC116 clones. AKT phosphorylates BAD and inhibits its association with Bcl-xL, there by inhibiting apoptosis. In CYC116 clones both AKT and Bcl-xL are overexpressed. Previously it was shown that CYC116 induced some CYP1A in human hepatocytes (10). Pathway analysis showed interaction of up-regulated CYP1A1 with retinoic acid derivatives. These interactions indicate the possibility of CYC116 as a substrate for CYP1A and related genes. Moreover some pathways relevant to Aurora kinase inhibition were also affected including cell cycle regulation of G1/S transition, initiation of mitosis, spindle assembly and chromosome separation, and DNA damage. Role of other observed affected pathways in the context of CYC116 and ZM447439-induced drug resistance are not clear. Taken together, the results suggest that tumor resistance towards CYC116 and ZM447439 is not mediated by one gene or one pathway, rather it is multifactorial. The drug resistance gene expression signatures specific to CYC116, could be used in the clinic to predict therapeutic response. As the mode of action of CYC116 and ZM447439 are similar to other AKIs, supported by the fact that CYC116 and ZM447439 resistant clones were cross-resistant to AZD1152, VX680, and MLN8054, the genetic fingerprint we have identified may be useful to predict the response to AKIs in general. Interestingly, CYC116 did not induce AK mutations as a mechanism of induced resistance, which could be advantageous in the clinic to design combinations to overcome relatively less aggressive resistant mechanisms. Moreover, our modeling studies showed that CYC116 can potentially inhibit the Aurora kinase with ZM447439-induced mutations that are likely to occur in the clinic. This is further supported by the fact that the cell lines harboring these mutations are significantly less cross-resistant to CYC116 (Table 1).

In conclusion, we have i) described mechanisms underlying resistance to novel AKI CYC116 in comparison to model compound ZM447439 and other clinically relevant AKIs; ii) identified and validated gene signatures associated with response to AKIs potentially usable for patient stratification; iii) showed that CYC116 is fully or partially active in mutant forms of Aurora B kinase associated with resistance against quinazoline class of AKIs; iv) and showed that resistance phenotype can be reversed, at least partially, using genetic or pharmacological inhibition of Bcl-xL protein. Thus, the CYC116 in combination with Bcl-xL inhibitors might be potentially useful to overcome or even prevent the emergence of resistance against AKIs, and the Bcl-xL inhibitors might be highly active in tumors resistant or refractory to synthetic AKIs.

## Acknowledgements

**Grant Support:** This study was supported by the Grant Agency of the Czech Republic (301/08/1649 and 303/09/H048) and Internal Grant Agency of the Palacky University (IGA UP LF 2011 018). Infrastructural part of this project (Institute of Molecular and Translational Medicine) was supported by the Operational Programme Research and Development for Innovations (project CZ.1.05/2.1.00/01.0030). The modeling part of the work was supported by a part of research project no. RVO: 61388963 of the IOCB, Academy of Sciences of the Czech Republic and was supported by Czech Science Foundation [P208/12/G016]. This work was also supported by the Operational Program Research and Development for Innovations - European Science Fund (CZ.1.05/2.1.00/03.0058). The support of Praemium Academiae of the Academy of Sciences of the Czech Republic awarded to P.H. in 2007 is also acknowledged. The financial support from the Gilead Sciences and IOCB Research Center, Prague, is also acknowledged. We are thankful to Cancer Research Foundation in Olomouc for fellowship awarded to Madhu Kollareddy.

