## Supplementary information for "Identification and characterization of drug resistance mechanisms in cancer cells against Aurora kinase inhibitors CYC116 and ZM447439"

**Supplemental Figures**

**Figure S1.** Sequenograms of wild-type and mutated Aurora B. Aurora B sequencing revealed three novel point mutations in ZM447439-resistant clones. cDNA sequences of Aurora B in parent cell lines (upper panel) vs. ZM447439-resistant clones (lower panel) at specific mutation site is presented in the sequenograms.

**Figure S2.** Unsupervised clustering of resistant clones and parent cell lines based on global gene expression patterns and heatmap created based on 50 genes, which mostly affect each of the first, second, and third component in PCA (total 138 genes, 12 genes overlap between the components of PCA). A, Dendrogram was created based on the global gene expression of all three clones (average of expression from three replicates) from all groups. Clones were clustered (unsupervised clustering) with respect to p53 background and compound used to generate resistant clones. This indicates that majority of the genes expression trends were common between the clones. B, 50 genes were selected, which mostly affect the first, second , and the third component (totally 138 genes, 12 genes overlap between the three components) in PCA, to create the heatmap. Here also the clones were clustered with respect to p53 background and compound used to generate resistant clones.

**Figure S3.** Correlation plot of 28 selected genes showing similar gene expression patterns between the microarray analysis and qRT-PCR analysis. Significant correlation (R=0857, p<0.000006) was achieved between the assay formats

**Figure S4.** GeneGo pathway analysis of top commonly affected pathway in each group of resistant clones. Top common pathway for each group of resistant clones is shown. A, Apoptotis and survival BAD phosphorylation pathway in HCT116 p53+/+:CYC116 clones. B, Retinol metabolism pathway in HCT116 p63-/-:CYC116 clones. C, Immune-response-NK2D signaling in HCT116 p53+/+:ZM447439 clones. D, Cell cycle regulation of mitosis in HCT116 p53-/-:ZM447439 clones.

**Figure S5.** Area under curves graphs for most significant individual genes or combination of five together. Top panel shows the significance of comparison of ROC (receiver operating characteristic) with combination model. The graph shows AUC (area under curve) for each gene and combinations.

**Figure S6.** Biomarker modulations in resistant clones in comparison to parent cell lines. A & B are the dot plots of parent DMSO controls. No inhibition of phospho-histone H3 can be seen. Here also from each group one resistant clone is shown. C, D, E, F, represents pH3 levels in resistant clones. G, H, I, J are parent controls treated with either CYC116 or ZM447439 for 24 h. Flow cytometry based assay for each sample was done in 3 biological replicates. K: The same profile can be noticed from western blotting (C=DMSO, C: 24 h = parent cell line treated with either CYC116 or ZM447439 for 24 h). Tubulin was used as loading control as shown in Fig. 1 (same lysates were used to determine phospho histone H3 (ser10) levels also)

**Figure S7.** Western blot showing protein expression levels of Aurora A and B in all CYC116 and ZM447439 resistant clones in comparison to DMSO controls. Tubulin was used as loading control as shown in Fig. 1 (same lysates were used for Aurora A and B expression studies also)

**Figure S8.** Tumorigenecity of CYC116 resistant clones. All CYC116 resistant clones were subcutaneously xenografted on both right and left side flanks of SCID mice. On each side 5-10 x 10^6^ cells were injected. The tumor volumes were measured from all three replicates. As shown in the graph, the tumor volumes of the xenograft increased significantly in a time dependent manner, indicating the proliferative and tumorigenic potential of the CYC116 resistant clones.

Figure S1

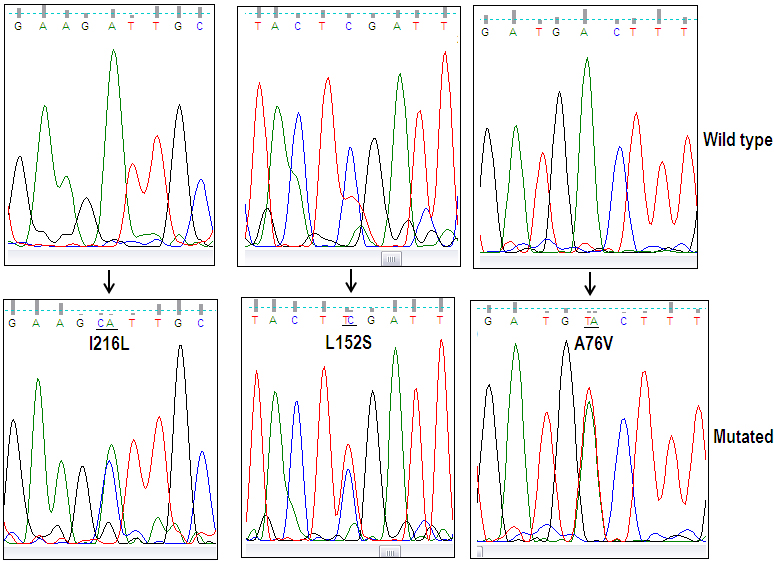

Figure S2

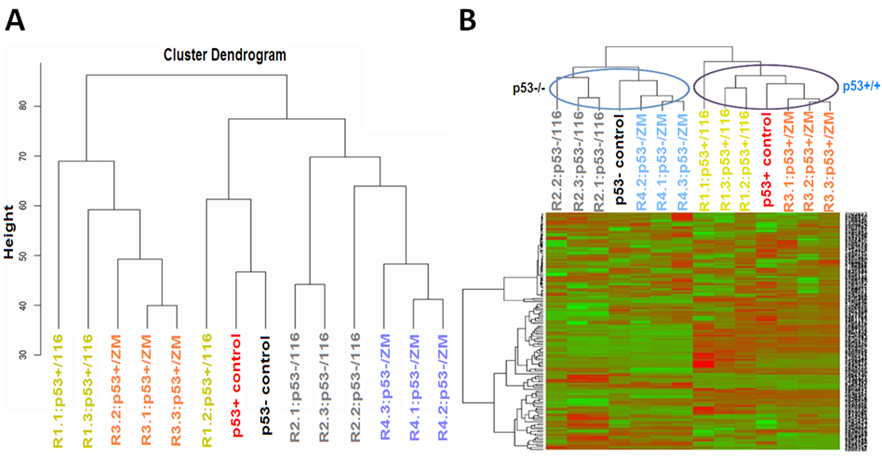

Figure S3

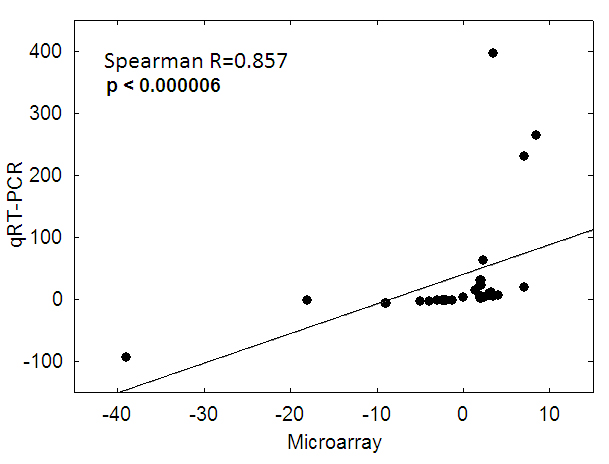

Figure S4A, B, C, D

S4A

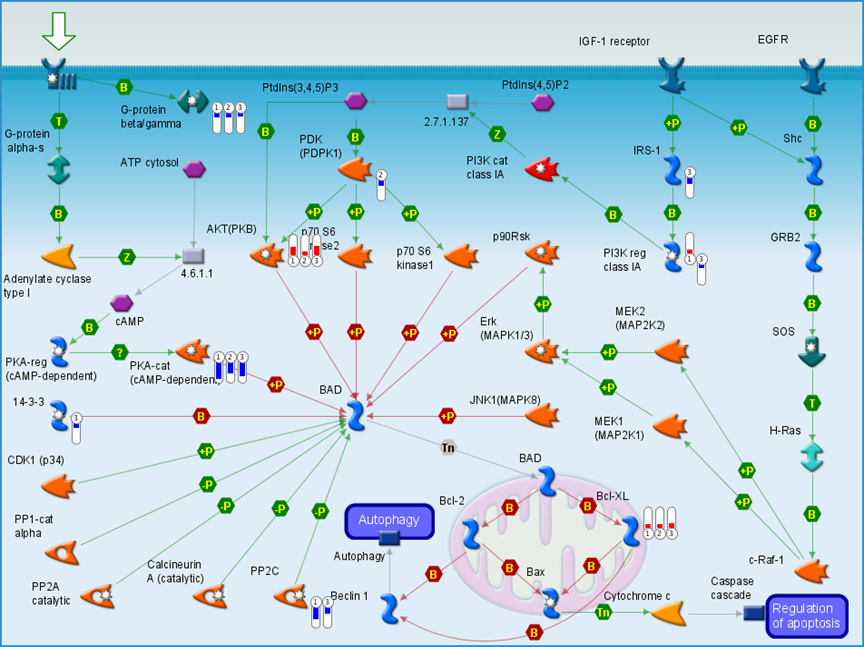

S4B

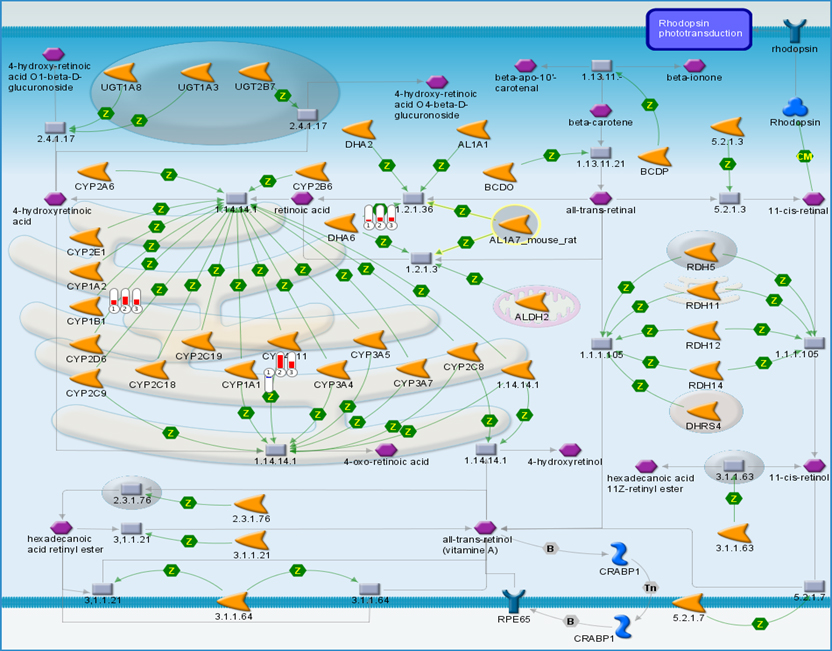

S4C

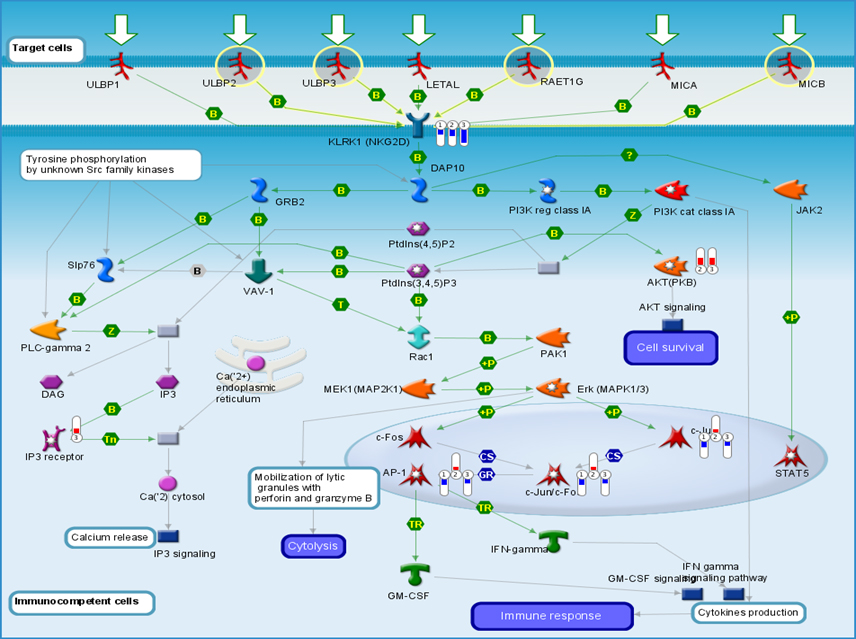

S4D

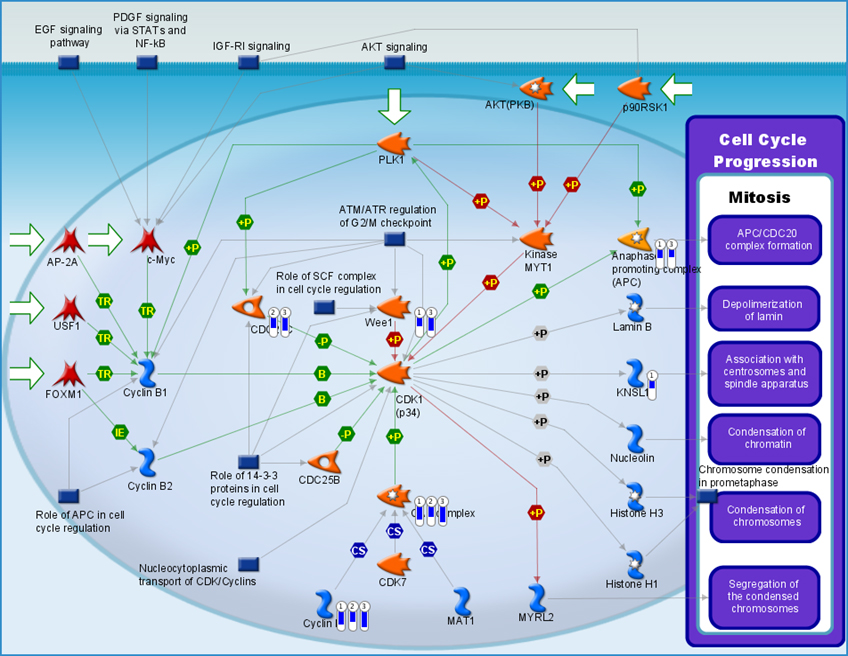

Figure S5

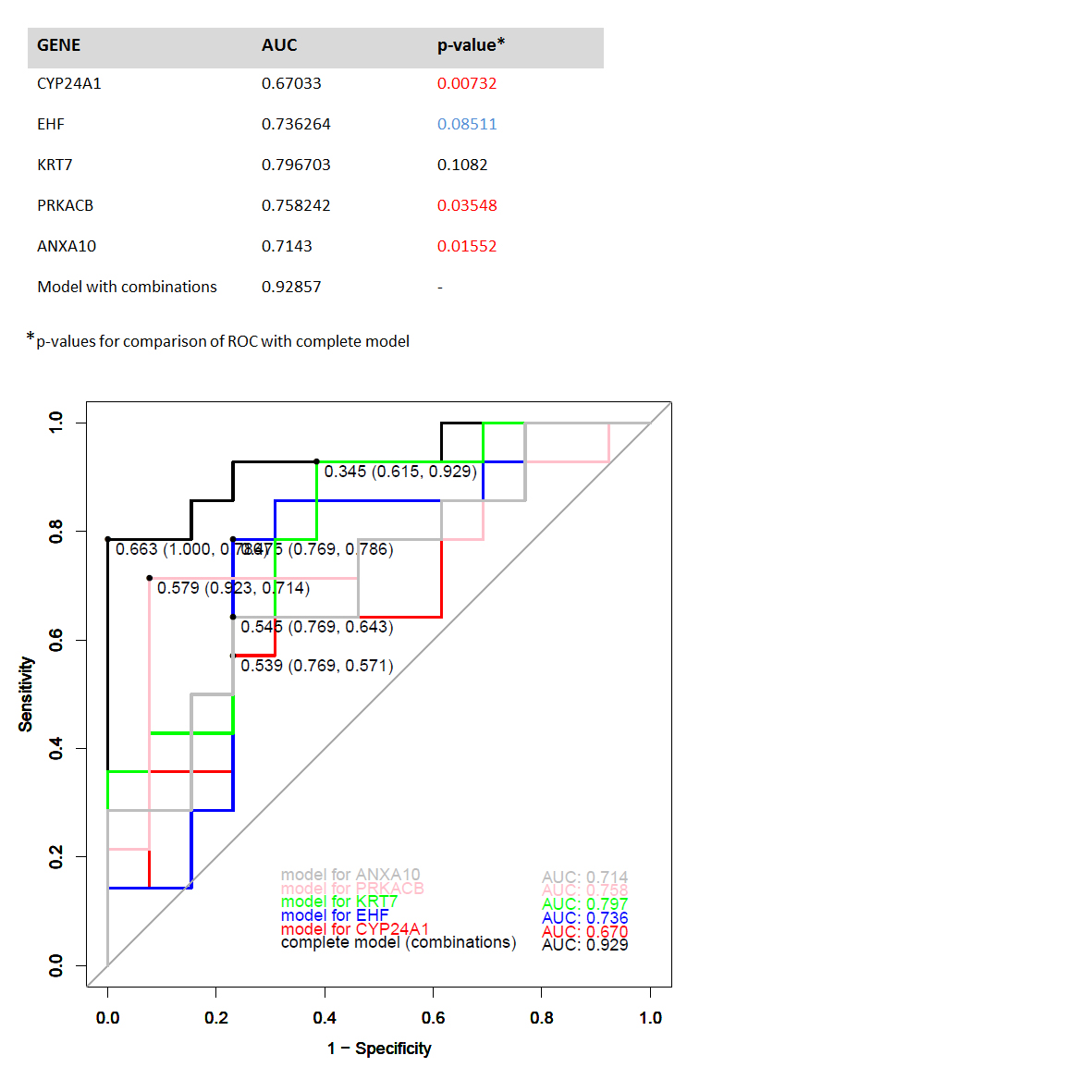

Figure S6

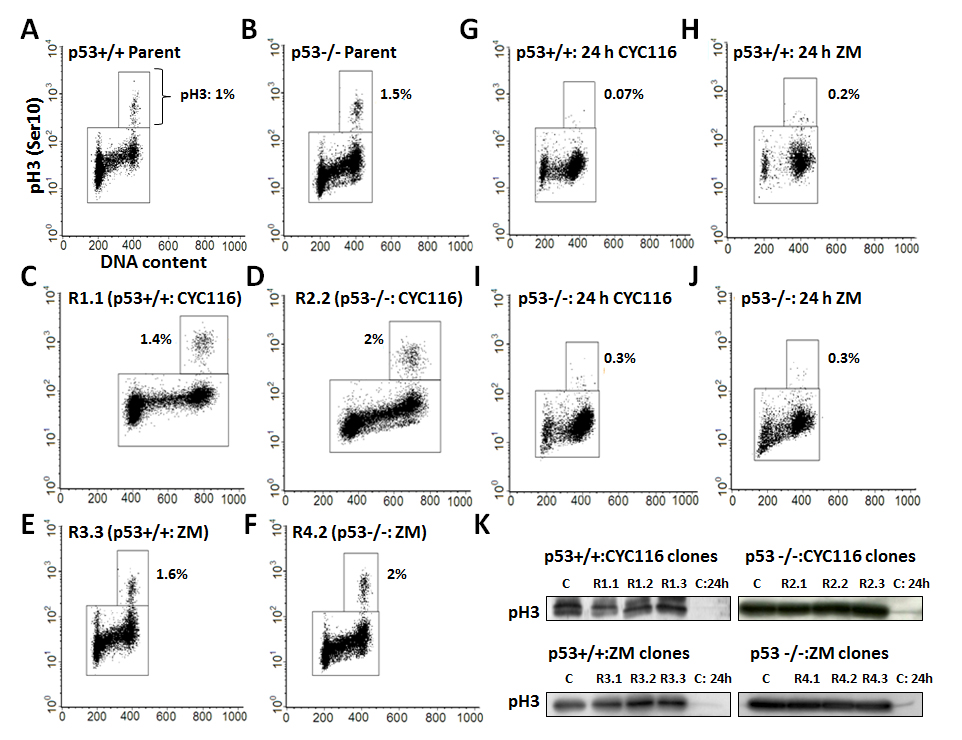

Figure S7

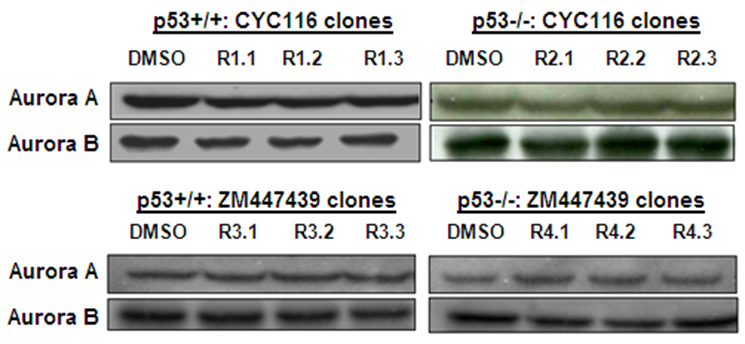

Figure S8

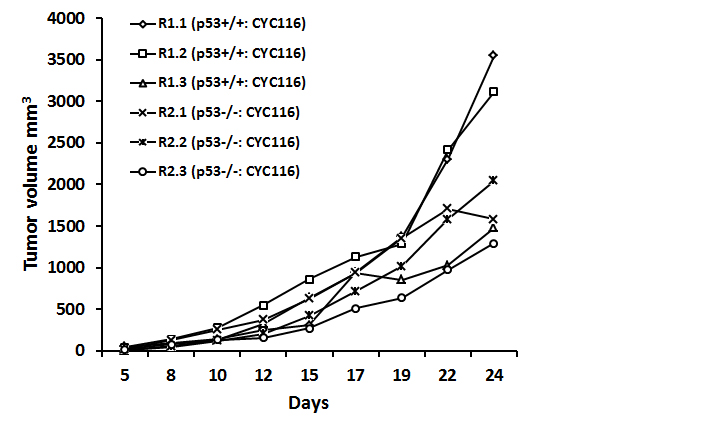

**Supplemental table legends**

**Supplemental Table S1.** Primers used for DNA sequencing of Aurora kinases. ^a^cDNA: complementary DNA, ^b^gDNA: Genomic DNA. Primer sequences used for Aurora A, B, and C kinases sequencing

**Supplemental Table S2.** Primers and thermal profiles used for qRT-PCR.

**Supplemetal Table S3.** MDR and sensitivity profiles of CYC116 and ZM447439 resistant clones. All IC50 values in the table are in micrograms (except bortezomib: µM), calculated from 3 independent replicates, each performed in two technical replicates. The SD values for the above data are in the range ± 0.000007 - ±4. IC50 values were also shown for p53+/+ and p53-/- parent cell lines. From each group two clones were selected to verify multidrug resistant (MDR) phenomenon using 13 approved anticancer agents.

**Supplemental Table S4.** Binding free energies ΔG_int_^w^ (kcal/mol) of ZM447439 and CYC116 with the wild type and mutated Aurora B proteins.

**Supplemental Table S5A.** Top 100 common differentially expressed genes (Cumulative p-value <0.001) and corresponding copy number changes in HCT116:CYC116 group. ^a^Chr. - Chromosome, ^b^logFC – Fold change, ^c^Amp. – Amplification, ^d^Del. – Deletion, ^e^Nd - No description. For some genes, identity number is presented more than once as respective Affymetrix probe binds to one more than one location of the genome having same recognition sequence. The same Gene IDs represented more than once, have unique ENSEMBL IDs.

**Supplemental Table S5B.** Top 100 common differentially expressed genes (Cumulative p-value <0.001) and corresponding copy number changes in HCT116 p53-/-:CYC116 group.

**Supplemental Table S5C.** Top 100 common differentially expressed genes (Cumulative p-value <0.001) and corresponding copy number changes in HCT116:ZM447439 group. **^a^**fg: family gene.

**Supplemental Table S5D.** Top 100 common differentially expressed genes (Cumulative p-value <0.001) and corresponding copy number changes in HCT116 p53-/-:ZM447439 group.

**Supplemental Table S6.** Totally 28 genes from microarray data (p<0.001) were validated by qRT-PCR. Nearly 100% correspondence in expression patterns can be noticed. Positive value indicate up-regulation and negative values represent down-regulation. Values from microarray data are represented as fold changes in comparison to control. ^a^NC=No change in expression.

**Supplemental Table S7.** Common and differential affected pathways based on p53 background of CYC116 or ZM447439 resistant clones.

**Supplemental Table S1**

| **Aurora Kinase** | **Reverse** | **Forward** |
| --- | --- | --- |
| Aurora A (cDNA^a^) | **(1)** TCAGGATTATTTTCAGGTGCCG | **(2)** TCCTTCAAATTCTTCCCAGCG |
|  | **(3)** AGAGAGTGGTCCTCCTGG | (**3)**TGGCAAGAGAAAAGCAAAGCAAG |
|  |  | **(4)** TGCCCTGTCTTACTGTCATTCG |
|  |  | **(5)** GCAAACACATACCAAGAGACC |
| Aurora B (cDNA) | **(1)** GTAGAGACGCAGGATGTTGG | **(2)** TTGATGACTTTGAGATTGGGCG |
|  |  | **(3)** GGAGGAAGACAATGTGTGGC |
| Aurora B (gDNA^b^) Exon2 |  | **(1)** TAGATCAGAGGGTCCGTTGG |
| Aurora C (gDNA)  Exons 1-4 | **(1)** AAAGAAGAGCGTTGGGGAGG |  |
|  | **(2)** CGGAGGGAAAGTCAGGGATG | **(3)** GAACTACTGATAGGGCTGGG |
|  | **(4)** TTCTCAGAAGGCAATGCGGA |  |
| Aurora C (cDNA)  Exons 5-7 | **(1)** GACAAATGAGGTGGCAGAGC | **(2)** AGCGAGAAATTAGATGAACAGCG |

**Supplemental Table S2**

| **Gene Symbol** | **Forward primer** | **Reverse primer** | **Thermal profiles** |
| --- | --- | --- | --- |
| CYP24A1 | CTGGGATCCAAGGCATTCTA | ATGGTGCTGACACAGGTGAA | 62° C/15 s |
| BCL2L1 | CTGGCTCCCATGACCATACT | GCTGAGGCCATAAACAGCTC | 62° C/15 s |
| GJC1 | ATGGTGTTACAGGCCTTTGC | GAGTCTCGAATGGTCCCAAA | 62° C/15 s |
| NCAM1 | TGAGTGGAGAGCAGTTGGTG | TTGGCATCATACCACTTGGA | 62° C/15 s |
| KLK5 | CTCGTGTCCTGGGGAGATTA | TGAACTTGCAGAGGTTCGTG | 62° C/15 s |
| KRT7 | GATGCTGCCTACATGAGCAA | TGAGGGTCCTGAGGAAGTTG | 62° C/15 s |
| LCN2 | CAAGGAGCTGACTTCGGAAC | GACAGGGAAGACGATGTGGT | 64° C/15 s |
| TNFAIP3 | ATGCACCGATACACACTGGA | GGATGATCTCCCGAAACTGA | 62° C/15 s |
| KRT13 | CGAGAGCCTGAATGAAGAGC | CGACCACCTGGTTGCTAAAT | 62° C/15 s |
| PPAP2B | AAATGACGCTGTGCTCTGTG | ACCGCGACTTCTTCAGGTAA | 62° C/15 s |
| TBX3 | GGGACATCGAACCTCAAAGA | CCATGCTCCTCTTTGCTCTC | 62° C/15 s |
| SERINC2 | CGTGTGGGTGAAGATCTGTG | CAGGGTCCACAGGTAGAGGA | 66° C/15 s |
| HOXB5 | AGGGCCCAAAGCTTGTAAAT | GCATCCACTCGCTCACTACA | 62° C/15 s |
| ANXA10 | GTCCTATGGGAAGCCTGTCA | GCTCTTGTTGCACAGGATCA | 60° C/15 s |
| CYP1A1 | GACAGATCCCATCTGCCCTA | CGAAGGAAGAGTGTCGGAAG | 62° C/15 s |
| PRKACB | GAGACCGTCCTTGTTGAAGC | ACGGGATGATGGCAATAAAG | 60° C/15 s |
| A4GALT | GACCACTACAACGGCTGGAT | CGGATGGAACACCACTTCTT | 62° C/15 s |
| ARHGAP29 | CATGGCAGCTGAATCTTTGA | AGCCAGATGACAGGAGCCTA | 62° C/15 s |
| NRP1 | CAAGGCGAAGTCTTTTGAGG | TCTCGGGGTAGATCCTGATG | 64° C/15 s |
| KLRK1 | GCCACAGCAGAGAGACACAG | CCCATTAAAAGTGGCAGCAT | 62° C/15 s |
| MID1 | ACCCAACATCAAGCAGAACC | GGCCTTGACCATGAAGATGT | 64° C/15 s |
| EHF | AGGTGATGCATCCTCCTCAC | AATGTTCACCTCCCTTGACG | 62° C/15 s |
| SEMA3A | TGCCAAGGCTGAAATTATCC | GCCAAGCCATTGAAAGTGAT | 62° C/15 s |
| PLK4 | TCCTTTTCCATTTGCAGACC | GCAGATTCCCAAACCACTGT | 64° C/15 s |
| INPP4B | GTGCTCCTTCAGGAACTTGC | AGTGCTTGGCTGAAGACGAT | 64° C/15 s |
| CAMK2D | CAGTACATGGATGGCAGTGG | TGCCACACACGAGTCTCTTC | 62° C/15 s |
| BDNF | CAAGGGGACCCATAGGAAAT | GAGCAAGGCACCTTCAAGTC | 62° C/15 s |
| TSPAN1 | CCTTTCTGCTCCAGACTTGG | AAGTCAGGCATCGCCTAAAA | 62° C/15 s |
| GAPDH | GAAGATGGTGATGGGATTTC | GAAGGTGAAGGTCGGAGT | 60° C/30 s |

**Supplemetal Table S3**

| **Drug** | **HCT116** | **R1.2:116**  **(p53 WT)** | **R1.3:116**  **(p53WT)** | **[**  **Drug** | **[**  **HCT116**  **p53-/-** | **R2.1:116**  **(p53-/-)** | **R2.2:116**  **(p53-/-)** |
| --- | --- | --- | --- | --- | --- | --- | --- |
| **Etoposide** | 1.3 | 45 (34) | 71 (53) | **Etoposide** | 1.84 | 14 (8) | 8.5 (5) |
| **Gemcitabine** | 0.08 | 2.3 (29) | 0.3 (4) | **Act-D** | 0.0015 | 0.004 (3) | 0.005 (3) |
| **Daunorubicin** | 0.03 | 0.33 (11) | 0.4 (16) | **Carboplatin** | 9.9 | 27 (3) | 26.3 (3) |
| **Act-D** | 0.0006 | 0.0027 (5) | 0.003 (6) | **Paclitaxel** | 0.004 | 0.008 (2) | 0.008 (2) |
| **Topotecan** | 0.01 | 0.045 (4.5) | 0.17 (17) | **Cladribine** | 7.6 | 16.5 (2) | 8 (1) |
| **Bortezomib** | 0.03 | 0.12 (4) | 0.3 (10) | **Cisplatin** | 2.1 | 3.35 (1.6) | 4.3 (2) |
| **Paclitaxel** | 0.0015 | 0.0055 (4) | 0.007 (5) | **Topotecan** | 0.155 | 0.21 (1.4) | 0.115 (0.7) |
| **Cisplatin** | 0.9 | 3.3 (4) | 2.37 (3) | **5-Flurouracil** | 1.2 | 1.5 (1.3) | 1 (0.8) |
| **Carboplatin** | 7 | 22 (3) | 12 (2) | **Oxaloplatin** | 3.3 | 2.05 (0.6) | 2.46 (0.7) |
| **Oxaloplatin** | 0.9 | 2.8 (3) | 1.2 (1) | **Gemcitabine** | 1.1 | 0.64 (0.6) | 0.63 (0.6) |
| **Doxorubicin** | 0.1 | 0.17 (1.7) | 0.2 (2) | **Doxorubicin** | 0.6 | 0.2 (0.3) | 0.32 (0.5) |
| **Cladribine** | 3.7 | 4.7 (1) | 4 (1) | **Daunorubicin** | 0.7 | 0.225(0.3) | 0.3 (0.4) |
| **5-Flurouracil** | 0.9 | 0.78 (0.8) | 1 (1) | **Bortezomib** | 0.28 | 0.04 (0.1) | 0.047 (0.2) |
|  |  | **[**  **R3.1:ZM**  **(p53 WT)** | **R3.2:ZM (p53WT)** |  |  | **R4.2:ZM**  **(p53-/-)** | **R4.3:ZM**  **(p53-/-)** |
| **Etoposide** |  | 40 (30) | 1.72 (1.3) | **Etoposide** |  | 39 (21) | 37 (20) |
| **Daunorubicin** |  | 0.25 (10) | 0.05 (2) | **Topotecan** |  | 0.4 (2.6) | 0.4 (2.6) |
| **Topotecan** |  | 0.04 (4) | 0.016 (2) | **5-Flurouracil** |  | 2 (1.6) | 2 (1.6) |
| **Carboplatin** |  | 25 (3.6) | 14 (2) | **Cladribine** |  | 16.5 (2) | 17 (2) |
| **Taxol** |  | 0.005 (3) | 0.001 (6) | **Oxaloplatin** |  | 5.9 (2) | 7 (2) |
| **Cisplatin** |  | 2.4 (3) | 2.42 (3) | **Bortezomib** |  | 0.5 (2) | 0.44 (1.6) |
| **Oxaloplatin** |  | 2.9 (3) | 1 (1) | **Paclitaxel** |  | 0.006 (1.5) | 0.012 (3) |
| **Act-D** |  | 0.002 (3) | 0.0009 (2) | **Act-D** |  | 0.002 (1.3) | 0.002 (1.3) |
| **Doxorubicin** |  | 0.19 (2) | 0.0355(0.4) | **Carboplatin** |  | 12 (1.2) | 14 (1.4) |
| **Cladribine** |  | 6.3 (2) | 2.3 (0.6) | **Cisplatin** |  | 2.27 (1) | 3.2 (1.5) |
| **5-Flurouracil** |  | 1.4 (1.5) | 0.46 (0.5) | **Gemcitabine** |  | 0.35 (0.3) | 0.3 (0.3) |
| **Gemcitabine** |  | 0.06 (0.8) | 0.026 (0.3) | **Doxorubicin** |  | 0.1 (0.15) | 0.2 (0.2) |
| **Bortezomib** |  | 0.006 (0.2) | 0.03 (1) | **Daunorubicin** |  | 0.1 (0.14) | 0.26 (0.4) |

**Supplemental Table S4**

| **Protein** | **ΔG_int_^w^ (kcal/mol)** | |
| --- | --- | --- |
|  | **ZM447439** | **CYC116** |
| Wild type Aurora B | -79.9 | -68.5 |
| I216L | -80.3 | -67.5 |
| L152S  N76V | -76.4 | -66.0 |
| L152S | -77.9 | -66.0 |
| G160E | - | -60.19 |
| G160V | - | -71.4 |
| Y156H | - | -69.8 |

**Supplemental Table S5A**

| **Gene ID** | **Gene Symbol** | **Chr.^a^** | **R1.1 logFC^b^** | **R1.2 logFC** | **R1.3**  **logFC** | **logFC Mean** | **R1.1**  **Copy No.** | **R1.2**  **Copy No.** | **R1.3**  **Copy No.** |
| --- | --- | --- | --- | --- | --- | --- | --- | --- | --- |
| 8067140 | CYP24A1 | 20 | -6.68 | -3 | -6.17 | -4.99 |  |  |  |
| 8047738 | NRP2 | 2 | 4.04 | 0.82 | 0.89 | 1.435 |  |  |  |
| 8065569 | BCL2L1 | 20 | 0.80 | 0.71 | 0.91 | 2.000 | Amp.^c^ | Amp. |  |
| 8047763 | Nd^e^ | 2 | 4.03 | 0.45 | 1.25 | 1.309 |  |  |  |
| 7964927 | TSPAN8 | 12 | 4.64 | 4.48 | 0.96 | 2.711 |  |  |  |
| 7944931 | SLC37A2 | 11 | 3.79 | 1.09 | 0.66 | 1.396 | Amp. | Amp. |  |
| 8016094 | GJC1 | 17 | -3.63 | -0.44 | -1.8 | -1.42 |  |  |  |
| 8152617 | HAS2 | 8 | -0.42 | 4.5 | 1.68 | 1.476 |  |  |  |
| 7961891 | BHLHE41 | 12 | 2.71 | -0.01 | 0.06 | 0.096 |  |  | Amp. |
| 7963614 | ITGB7 | 12 | 3.93 | 1.06 | 1.4 | 1.802 |  |  |  |
| 8101828 | TSPAN5 | 4 | -4.39 | -0.78 | -1 | -1.51 |  |  |  |
| 8150529 | DKK4 | 8 | -0.05 | -0.07 | 3.56 | 0.233 |  |  |  |
| 8070574 | TFF2 | 21 | 2.02 | -0.25 | 0.14 | 0.411 |  | Amp. | Amp. |
| 7935553 | LOXL4 | 10 | 3.21 | 0.04 | 0.77 | 0.447 |  |  |  |
| 7943892 | NCAM1 | 11 | 2.87 | -0.1 | 2.94 | 0.944 | Amp. | Amp. |  |
| 8038670 | KLK5 | 19 | 4.23 | 0.37 | 1.17 | 1.227 |  | Amp. |  |
| 7955613 | KRT7 | 12 | 3.71 | -0.22 | 1.29 | 1.018 |  |  |  |
| 8158167 | LCN2 | 9 | 5.3 | 1.71 | 2.22 | 2.723 |  | Amp. |  |
| 8122265 | TNFAIP3 | 6 | 2.36 | 0.65 | 3.11 | 1.686 |  |  |  |
| 8015323 | KRT13 | 17 | 5.5 | 0.72 | 1.37 | 1.755 |  | Amp. |  |
| 8020740 | DSG4 | 18 | 2.69 | 0.23 | -0.15 | 0.455 |  |  |  |
| 8123936 | NEDD9 | 6 | 2.47 | 0.03 | 0.27 | 0.262 |  |  | Del.^d^ |
| 8173261 | ZC4H2 | X | 0.3 | -0.05 | -1.82 | -0.29 |  |  |  |
| 8152606 | SNTB1 | 8 | 0.12 | 3.06 | 1.84 | 0.872 |  |  |  |
| 8016994 | RNF43 | 17 | -2.98 | 0.58 | 0.09 | -0.54 |  |  |  |
| 8168749 | SRPX2 | X | 2.71 | 0.28 | 0.78 | 0.84 |  |  |  |
| 8112615 | ENC1 | 5 | -2.39 | -1.49 | -2.01 | -1.93 |  |  |  |
| 7916493 | PPAP2B | 1 | 1.57 | 0.03 | 1.53 | 0.433 |  |  |  |
| 8081548 | PVRL3 | 3 | -3.43 | 0.18 | -1.01 | -0.85 |  |  |  |
| 8090180 | MUC13 | 3 | 1.12 | 3.14 | 0.16 | 0.818 |  | Amp. |  |
| 8135763 | WNT16 | 7 | -2.96 | 0.23 | -1.1 | -0.91 | Amp. | Amp. |  |
| 8138566 | IGF2BP3 | 7 | -3.22 | 0.26 | 0.31 | -0.64 | Amp. | Amp. |  |
| 8068633 | B3GALT5 | 21 | 2.21 | -0.16 | 0.27 | 0.454 |  |  | Amp. |
| 8140955 | CDK6 | 7 | -0.99 | 0.64 | 1.49 | 0.98 | Amp. |  |  |
| 8176174 | MPP1 | X | -1.87 | -0.06 | 0.06 | -0.19 |  |  |  |
| 8026468 | CYP4F12 | 19 | 2.49 | 0.62 | 0.85 | 1.095 |  |  |  |
| 8174598 | IL13RA2 | X | 3.4 | 0.58 | 0.35 | 0.881 |  |  |  |
| 8129677 | SGK1 | 6 | 2.27 | 1.61 | 1.44 | 1.739 |  |  |  |
| 8120043 | RUNX2 | 6 | 2.58 | 2.09 | 0.96 | 1.733 |  |  |  |
| 8038725 | KLK10 | 19 | 3.93 | 0.78 | 1.73 | 1.746 |  | Amp. |  |
| 8096116 | AGPAT9 | 4 | 2.68 | 1.14 | -0.58 | 1.211 |  |  |  |
| 8148548 | PSCA | 8 | 2.34 | -0.04 | 0.47 | 0.339 |  | Amp. |  |
| 8161964 | FRMD3 | 9 | 3.14 | 0.39 | 0.32 | 0.734 |  |  |  |
| 7970954 | DCLK1 | 13 | -0.44 | 2.21 | 3.21 | 1.463 |  |  | Del. |
| 7966690 | TBX3 | 12 | 2.29 | 1.39 | 1.58 | 1.714 |  |  | Amp. |
| 7899615 | SERINC2 | 1 | 2.44 | 2.13 | 2.37 | 2.312 |  | Amp. |  |
| 8049349 | UGT1A | 2 | 1.28 | -0.11 | 0.17 | 0.288 |  |  |  |
| 8106986 | RHOBTB3 | 5 | -1.64 | 0.15 | -3 | -0.91 |  |  |  |
| 8027748 | FXYD3 | 19 | 3.4 | 1.02 | 1.88 | 1.868 |  |  |  |
| 7973433 | DHRS2 | 14 | 0.45 | 0.87 | 2.2 | 0.95 | Del. | Del. |  |
| 8101675 | ABCG2 | 4 | 2.87 | 1.01 | 0.27 | 0.922 |  |  |  |
| 8151730 | CALB1 | 8 | 3.44 | 0.8 | 1.74 | 1.683 |  |  |  |
| 7927215 | ALOX5 | 10 | 2.78 | 0.73 | 1.59 | 1.479 |  |  |  |
| 8045889 | TANC1 | 2 | 1.68 | 0.3 | 0.33 | 0.552 |  |  |  |
| 7925531 | AKT3 | 1 | 1.98 | 0.91 | 2.19 | 1.578 |  | Amp. |  |
| 8098441 | ODZ3 | 4 | 1.57 | 0.28 | 1.61 | 0.896 |  |  | Del. |
| 8044574 | IL1RN | 2 | 1.81 | 0.1 | 0.24 | 0.354 |  | Del. |  |
| 8038683 | KLK6 | 19 | 3.25 | 0.93 | 0.87 | 1.381 |  | Amp. |  |
| 7922773 | NCF2 | 1 | 1.59 | 0.09 | 0.65 | 0.454 |  |  |  |
| 8068100 | NCRNA00189 | 21 | 0.11 | 0.29 | 1.35 | 0.347 |  |  | Amp. |
| 8037205 | CEACAM1 | 19 | 3.05 | 0.75 | 1.64 | 1.556 |  | Amp. |  |
| 7918657 | PTPN22 | 1 | 3.67 | 1.53 | 0.72 | 1.591 |  |  |  |
| 8098263 | PALLD | 4 | -1.96 | -1.72 | -2.27 | -1.97 |  |  | Del. |
| 8053417 | CAPG | 2 | 1.43 | -0.7 | -0.23 | 0.616 |  | Amp. |  |
| 8016457 | HOXB5 | 17 | 1.49 | 1.97 | 2.44 | 1.927 |  |  |  |
| 8067055 | ATP9A | 20 | 1.07 | 0.04 | -0.64 | 0.301 |  |  |  |
| 7902104 | PDE4B | 1 | -2.32 | -0.11 | -2.07 | -0.8 |  |  |  |
| 8077899 | PPARG | 3 | 2.26 | 0.56 | 0.56 | 0.89 |  |  |  |
| 8015016 | TNS4 | 17 | 0.52 | 0.83 | 1.68 | 0.895 |  |  |  |
| 7915472 | SLC2A1 | 1 | -1.73 | 0.8 | 1.04 | 1.13 |  |  |  |
| 8095728 | EREG | 4 | -1.52 | 0.1 | -3.87 | -0.83 |  |  |  |
| 7923958 | C1orf116 | 1 | 2.01 | 0.54 | 0.82 | 0.96 |  |  |  |
| 7955694 | IGFBP6 | 12 | 2.27 | 1.12 | 1.5 | 1.56 |  |  |  |
| 8112803 | LHFPL2 | 5 | 1.39 | 0.1 | -0.15 | 0.273 |  |  |  |
| 8033780 | ZNF426 | 19 | -1.11 | 1.12 | -0.92 | -1.04 |  |  |  |
| 8016463 | HOXB6 | 17 | 1.53 | 2.06 | 2.45 | 1.979 |  |  |  |
| 7940643 | ASRGL1 | 11 | -1.35 | 0.56 | 0.01 | -0.2 |  | Amp. |  |
| 7961182 | KLRC2 | 12 | -3.17 | -0.99 | -1.91 | -1.82 |  |  | Amp. |
| 8038695 | KLK7 | 19 | 2.78 | 0.72 | 0.82 | 1.178 |  | Amp. |  |
| 7950534 | WNT11 | 11 | 2.45 | 0.77 | 0.45 | 0.951 | Amp. | Amp. |  |
| 7986214 | SLCO3A1 | 15 | 2.27 | 0.53 | 1.26 | 1.148 |  |  |  |
| 8098246 | ANXA10 | 4 | -0.19 | -1.75 | -1.4 | -0.77 |  |  |  |
| 7990391 | CYP1A1 | 15 | 2.51 | 1.14 | 0.91 | 1.374 |  |  |  |
| 7946781 | PLEKHA7 | 11 | 1.68 | 0.52 | 0.43 | 0.722 | Amp. | Amp. |  |
| 8070411 | C21orf88 | 21 | 1.43 | -0.21 | 0.11 | 0.32 |  |  | Amp. |
| 7920128 | S100A11 | 1 | 1.24 | 0.69 | 1.6 | 1.108 |  | Amp. |  |
| 7902594 | PRKACB | 1 | -3.7 | -2.59 | -3.14 | -3.11 |  |  |  |
| 7957023 | LYZ | 12 | 3.63 | 0.7 | 1.24 | 1.466 |  |  |  |
| 8150509 | PLAT | 8 | 1.92 | -0.61 | 0.77 | 0.968 |  |  |  |
| 7920285 | S100A2 | 1 | 1.43 | -0.12 | -7.87E-05 | 0.024 |  | Amp. |  |
| 7976425 | OTUB2 | 14 | 1.56 | 0.69 | 0.81 | 0.957 | Del. |  |  |
| 8122146 | Nd | 6 | -2.21 | 0.89 | 0.2 | -0.74 |  |  |  |
| 8042993 | CTNNA2 | 2 | 1.1 | -0.03 | 0.33 | 0.227 |  |  |  |
| 8076497 | A4GALT | 22 | 1.39 | 1 | 2.15 | 1.439 |  | Amp. |  |
| 8073068 | APOBEC3C | 22 | 1.82 | 1.35 | 1.77 | 1.633 |  | Amp. |  |
| 7917850 | ARHGAP29 | 1 | -4.1 | -1.54 | -1.73 | -2.22 |  |  |  |
| 7938035 | TRIM22 | 11 | 1.04 | 1.76 | 0.49 | 0.964 | Amp. |  |  |
| 7932985 | NRP1 | 10 | 2.95 | -0.18 | 0.18 | 0.458 |  |  |  |
| 7961151 | KLRK1 | 12 | -4.33 | -0.91 | -2.15 | -2.04 |  |  | Amp. |
| 7963333 | KRT80 | 12 | 1.51 | -0.15 | -0.03 | 0.199 |  |  |  |

**Supplemental Table S5B**

| **Gene ID** | **Gene symbol** | **Chr.** | **R2.1 logFC** | **R2.2 logFC** | **R2.3 logFC** | **logFC Mean** | **R2.1**  **Copy No.** | **R2.2 Copy No.** | **R2.3**  **Copy No.** |
| --- | --- | --- | --- | --- | --- | --- | --- | --- | --- |
| 8135763 | WNT16 | 7 | -0.6 | -3.9 | -0.38 | -0.95 |  |  |  |
| 7906954 | PBX1 | 1 | 1.38 | 4.11 | 1.36 | 1.98 |  |  |  |
| 8140955 | CDK6 | 7 | -2.05 | 1.29 | -1.69 | -1.65 |  | Amp. |  |
| 8171297 | MID1 | X | -3.99 | -4 | -4.66 | -4.19 | Del. | Del. | Del. |
| 7939314 | EHF | 11 | 5.37 | 1.13 | 4.74 | 3.07 |  |  |  |
| 8013384 | ALDH3A1 | 17 | 0.5 | 3.72 | 0.24 | 0.76 |  |  | Del. |
| 8046726 | SSFA2 | 2 | -0.47 | -2.1 | -0.51 | -0.8 | Del. |  | Del. |
| 8152376 | CSMD3 | 8 | -0.3 | 1.67 | -0.12 | 0.39 |  | Del. |  |
| 8067140 | CYP24A1 | 20 | -5.54 | -3.7 | -5.79 | -4.92 |  |  |  |
| 8140468 | PION | 7 | 4.09 | -0.2 | 3.51 | 1.44 |  |  |  |
| 7895417 | SEPT2 | 2 | -1.83 | -0.1 | -2.04 | -0.6 |  |  |  |
| 8106727 | ATP6AP1L | 5 | 2.49 | -0.2 | 2.26 | 1.01 | Amp. | Amp. | Amp. |
| 7951686 | IL18 | 11 | 0.6 | -1.7 | 0.58 | -0.84 | Amp. | Amp. | Amp. |
| 8148309 | Nd | 8 | -1.39 | -1.7 | -1.18 | -1.42 |  | Del. |  |
| 8140668 | SEMA3A | 7 | 0.48 | -2.5 | 0.56 | -0.87 |  |  |  |
| 8081548 | PVRL3 | 3 | -0.51 | -2.4 | -0.6 | -0.9 |  | Amp. |  |
| 7950810 | SYTL2 | 11 | 1.44 | -1.6 | 1.2 | 1.42 | Amp. | Amp. | Amp. |
| 7910915 | CHRM3 | 1 | -0.19 | 2.02 | 0.13 | 0.37 |  | Del. |  |
| 8038695 | KLK7 | 19 | 1.48 | 0.1 | 1.61 | 0.61 |  |  |  |
| 7917850 | ARHGAP29 | 1 | -1.95 | -3.9 | -1.25 | -2.11 |  |  |  |
| 8113761 | ZNF608 | 5 | -1 | -1.7 | -0.98 | -1.19 | Amp. | Amp. | Amp. |
| 8076497 | A4GALT | 22 | 0.89 | 1.68 | 1.1 | 1.18 |  |  |  |
| 8122634 | SAMD5 | 6 | 2 | -0.3 | 1.6 | 1 |  |  |  |
| 7957298 | NAV3 | 12 | -0.04 | -2 | 0.11 | -0.21 |  |  |  |
| 8073096 | APOBEC3H | 22 | 1.71 | 0.86 | 1.84 | 1.39 |  |  |  |
| 8114119 | FSTL4 | 5 | 1.54 | 1.3 | 1.58 | 1.47 | Amp. |  | Amp. |
| 7958884 | OAS1 | 12 | 0.3 | 2.31 | 0.37 | 0.64 |  |  |  |
| 8121749 | GJA1 | 6 | 0.25 | -0 | 1.86 | 0.28 | Amp. | Amp. | Amp. |
| 8065569 | BCL2L1 | 20 | 0.53 | 0.86 | 0.41 | 1.5 |  |  |  |
| 7965941 | GLT8D2 | 12 | 0.94 | -0.8 | 0.88 | 0.86 |  |  |  |
| 8141066 | PON3 | 7 | -2.23 | -2.2 | -1.95 | -2.11 |  |  |  |
| 7906969 | Nd | 1 | 0.05 | 1.85 | 0.13 | 0.23 |  |  |  |
| 8023043 | PSTPIP2 | 18 | -0.01 | -1.3 | -0.24 | -0.15 | Amp. | Del. |  |
| 8097356 | PLK4 | 4 | -1.31 | -0.8 | -1.42 | -1.16 | Del. | Del. | Del. |
| 7962151 | DENND5B | 12 | 0.96 | 1.65 | 0.86 | 1.11 |  |  |  |
| 7932744 | ARMC4 | 10 | -0.38 | -1.9 | -0.33 | -0.62 |  |  |  |
| 7934161 | PRF1 | 10 | -2.9 | -2.2 | -2.8 | -2.63 | Amp. | Amp. | Amp. |
| 8127234 | DST | 6 | -1.27 | -2.2 | -1.36 | -1.57 | Amp. | Amp. | Amp. |
| 8084630 | Nd | 3 | 1.37 | 2.24 | 1.15 | 1.52 |  | Amp. |  |
| 8084630 | Nd | 3 | 1.37 | 2.24 | 1.15 | 1.52 |  | Amp. |  |
| 8084630 | Nd | 3 | 1.37 | 2.24 | 1.15 | 1.52 |  | Amp. |  |
| 8007446 | IFI35 | 17 | -0.46 | 2.23 | -0.45 | 0.77 |  |  |  |
| 8115490 | ADAM19 | 5 | 0.68 | -2 | 0.4 | -0.81 |  |  |  |
| 8082075 | DTX3L | 3 | -0.45 | 1.39 | -0.12 | 0.42 |  | Amp. |  |
| 8075310 | LIF | 22 | 1.3 | -0.2 | 1.35 | 0.66 |  |  |  |
| 8102950 | INPP4B | 4 | -0.68 | -2.7 | -1.01 | -1.23 | Del. | Del. | Del. |
| 8027748 | FXYD3 | 19 | 0.74 | 2.71 | 0.76 | 1.15 |  |  |  |
| 8065071 | FLRT3 | 20 | 0.34 | 1.64 | 0.21 | 0.49 |  |  |  |
| 8101828 | TSPAN5 | 4 | -1.08 | -2.8 | -1.11 | -1.49 | Del. | Del. | Del. |
| 8166747 | SYTL5 | X | 0.85 | -2.4 | 0.9 | -1.22 |  |  |  |
| 7990391 | CYP1A1 | 15 | 2.56 | 4.74 | 2.21 | 2.99 |  |  | Amp. |
| 8152506 | SAMD12 | 8 | 1.51 | 1.81 | 1.63 | 1.64 |  | Del. | Del. |
| 7927202 | ZNF22 | 10 | -2.48 | -2 | -2.29 | -2.23 | Amp. | Amp. | Amp. |
| 7902594 | PRKACB | 1 | -1.56 | -2 | -1.35 | -1.62 | Amp. | Amp. | Amp. |
| 8036318 | ZNF566 | 19 | -0.68 | 1.35 | -0.8 | -0.9 |  | Del. |  |
| 7935521 | AVPI1 | 10 | 1.08 | 1.17 | 1.19 | 1.15 | Amp. | Amp. | Amp. |
| 8022711 | DSC2 | 18 | -0.02 | -1.5 | -0.34 | -0.22 | Amp. | Del. | Amp. |
| 7932765 | MPP7 | 10 | -0.12 | -1.4 | -0.17 | -0.3 |  | Del. | Del. |
| 7957260 | GLIPR1 | 12 | -0.81 | -2.7 | -0.48 | -1.01 |  |  |  |
| 7916862 | WLS | 1 | 1.12 | -0.6 | 1.21 | 0.93 |  |  |  |
| 8102415 | CAMK2D | 4 | -0.66 | -1.7 | -0.77 | -0.95 | Del. | Del. | Del. |
| 8150830 | LYPLA1 | 8 | -1.23 | -1.1 | -1.07 | -1.12 | Del. | Del. | Del. |
| 8154135 | SLC1A1 | 9 | 1.03 | -1.8 | 0.97 | 1.21 | Amp. | Del. |  |
| 8148304 | TRIB1 | 8 | 0.03 | -0.9 | 0.23 | -0.18 |  | Del. |  |
| 8106743 | VCAN | 5 | 1.05 | -2.6 | 1.14 | -1.47 | Amp. | Amp. | Amp. |
| 8005029 | MAP2K4 | 17 | -1.2 | -0.6 | -1.38 | -1.01 | Del. |  | Del. |
| 8138566 | IGF2BP3 | 7 | -2.63 | -0.3 | -1.63 | -1.05 |  | Amp. |  |
| 8059716 | C2orf52 | 2 | 1.18 | 0.75 | 1.54 | 1.11 | Amp. | Amp. | Amp. |
| 8106986 | RHOBTB3 | 5 | -0.41 | -2 | -0.54 | -0.76 | Amp. | Amp. | Amp. |
| 8016094 | GJC1 | 17 | -2.55 | -1.9 | -2.36 | -2.24 | Amp. | Amp. |  |
| 8133018 | ZNF716 | 7 | 0.05 | 2.51 | 0.53 | 0.39 | Amp. | Amp. | Amp. |
| 8144758 | ZDHHC2 | 8 | 0.41 | -0.8 | 0.45 | 0.53 | Del. | Del. | Del. |
| 8129482 | SAMD3 | 6 | -0.07 | -1.2 | -0.1 | -0.2 | Amp. |  |  |
| 7917528 | Nd | 1 | -0.34 | 0.6 | -0.68 | -0.52 |  |  |  |
| 8100328 | USP46 | 4 | -0.84 | 0.11 | -0.85 | -0.43 | Del. | Amp. | Del. |
| 8047738 | NRP2 | 2 | -0.01 | 1.1 | 0.34 | 0.17 |  | Amp. |  |
| 7947230 | BDNF | 11 | -0.29 | -2.2 | -0.35 | -0.6 |  |  |  |
| 8081214 | GPR15 | 3 | 1.42 | -1.3 | 1.03 | 1.23 |  | Amp. |  |
| 8104107 | TRIML2 | 4 | -1.78 | -2 | -1.6 | -1.78 |  |  |  |
| 7892605 | SEPT2 | 2 | -1.5 | 0.12 | -1.33 | -0.62 |  |  |  |
| 8120176 | C6orf141 | 6 | 0.27 | -1.2 | 0.64 | -0.59 | Amp. | Amp. | Amp. |
| 7930498 | ACSL5 | 10 | -1.7 | -2 | -1.18 | -1.59 |  |  |  |
| 8060225 | HDLBP | 2 | -0.91 | -0.1 | -1.07 | -0.38 |  | Amp. | Amp. |
| 8152617 | HAS2 | 8 | 2.11 | 0.03 | 2.25 | 0.53 |  | Del. | Del. |
| 7935660 | DNMBP | 10 | -0.34 | -1.7 | -0.44 | -0.64 | Amp. |  |  |
| 8075910 | RAC2 | 22 | -0.01 | -1.2 | -0.06 | -0.08 |  |  |  |
| 8059345 | SCG2 | 2 | -1.05 | 0.23 | -1.16 | -0.65 |  | Amp. |  |
| 8081158 | ARL6 | 3 | -0.24 | 0.98 | -0.09 | 0.27 |  | Amp. |  |
| 8035095 | CYP4F11 | 19 | -1.87 | -0.7 | -2.06 | -1.36 |  |  | Amp. |
| 8160670 | AQP3 | 9 | 0.41 | 2.75 | 0.25 | 0.65 |  |  |  |
| 8141035 | SGCE | 7 | -1.18 | 0.39 | -0.64 | -0.67 |  |  |  |
| 8059111 | ABCB6 | 2 | -0.21 | 0.74 | -0.34 | 0.37 |  | Amp. | Amp. |
| 8059111 | ATG9A | 2 | -0.21 | 0.74 | -0.34 | 0.37 |  | Amp. | Amp. |
| 7988260 | FRMD5 | 15 | -1.5 | -1.7 | -1.38 | -1.52 | Amp. |  | Amp. |
| 7896498 | SEPT2 | 2 | -0.81 | -0 | -1.07 | -0.33 |  |  |  |
| 8017651 | SMURF2 | 17 | -1.08 | -1 | -1.14 | -1.06 | Amp. |  |  |
| 8146379 | UBE2V2 | 8 | -0.81 | -0.5 | -0.92 | -0.71 | Del. | Del. | Del. |
| 7993478 | ABCC1 | 16 | -0.2 | 1.12 | -0.17 | 0.33 |  | Amp. |  |
| 8017843 | SLC16A6 | 17 | 2.4 | -0.6 | 2.61 | 1.6 |  |  |  |
| 8112615 | ENC1 | 5 | 0.09 | -1.5 | 0.39 | -0.38 | Amp. | Amp. | Amp. |

**Supplemental Table S5C**

| **Gene ID** | **Gene symbol** | **Chr.** | **R3.1 logFC** | **R3.2 logFC** | **R3.3 logFC** | **logFC Mean** | **R3.1**  **Copy No.** | **R3.2 Copy No.** | **R3.3**  **Copy No.** |
| --- | --- | --- | --- | --- | --- | --- | --- | --- | --- |
| 8098441 | ODZ3 | 4 | 1.949 | 1.872 | 2.185 | 1.998 | Del. |  |  |
| 7932744 | ARMC4 | 10 | -2.59 | -2.67 | -2.52 | -2.59 | Amp. |  |  |
| 8144726 | TUSC3 | 8 | 1.872 | 2.211 | 2.602 | 2.209 | Amp. |  |  |
| 8098263 | PALLD | 4 | -2.18 | -2 | -1.99 | -2.05 | Amp. |  |  |
| 7989146 | MNS1 | 15 | -1.61 | -1.56 | -1.35 | -1.5 |  |  |  |
| 7894805 | Nd | 1 | -0.43 | -1.91 | -0.55 | -0.77 |  |  |  |
| 8021169 | LIPG | 18 | -1.03 | -1 | -1.22 | -1.08 |  |  |  |
| 8059854 | ARL4C | 2 | 1.866 | 0.953 | 1.152 | 1.27 |  |  |  |
| 7893924 | Nd | 5 | 4.604 | 6.218 | 5.593 | 5.43 |  |  |  |
| 7895294 | ILF2 | 1 | -1.37 | -1.33 | -0.49 | -0.96 |  |  |  |
| 8122176 | TCF21 | 6 | -1.22 | -0.97 | -1.06 | -1.08 |  |  |  |
| 7932765 | MPP7 | 10 | -2.08 | -2.28 | -2.2 | -2.18 | Amp. |  |  |
| 7895205 | Nd | 1 | 1.628 | 1.559 | 1.57 | 1.586 |  |  |  |
| 7894487 | Nd | 2 | -1.06 | -1.46 | -0.28 | -0.75 |  |  |  |
| 7893953 | Nd | 17 | 0.941 | 1.278 | 1.175 | 1.122 |  |  |  |
| 7975154 | NCRNA00238 | 14 | 1.573 | 0.154 | 0.215 | 0.373 | Del. |  |  |
| 7896206 | Nd | 14 | -0.39 | -1.42 | -0.71 | -0.73 |  |  |  |
| 7932733 | MKX | 10 | -1.76 | -1.68 | -1.75 | -1.73 | Amp. |  |  |
| 8152376 | CSMD3 | 8 | 1.521 | 1.813 | 1.934 | 1.747 | Amp. |  |  |
| 8112615 | ENC1 | 5 | -1.86 | -1.39 | -0.99 | -1.37 | Amp. |  |  |
| 8102328 | CFI | 4 | 0.822 | 0.178 | 0.071 | 0.218 | Del. |  |  |
| 8088952 | Nd | 3 | 1.552 | 0.431 | 0.654 | 0.759 |  |  |  |
| 7893175 | Nd | 19 | 1.829 | 1.995 | 1.755 | 1.857 |  |  |  |
| 8089467 | ZBED2 | 3 | -1.75 | -0.71 | -0.47 | -0.83 | Amp. | Amp. |  |
| 8013519 | Nd | 17 | 1.872 | 1.107 | 0.327 | 0.878 |  |  |  |
| 8013519 | Nd | 5 | 1.872 | 1.107 | 0.327 | 0.878 |  |  |  |
| 8003230 | Nd | 16 | 0.991 | 0.934 | 1.073 | 0.998 | Del. |  |  |
| 7899615 | SERINC2 | 1 | 0.523 | 1.289 | 1.146 | 0.917 | Del. |  |  |
| 7937335 | IFITM...fg^a^ | 11 | 2.179 | 0.229 | 0.228 | 0.484 | Del. |  |  |
| 7937335 | IFITM1 | 11 | 2.179 | 0.229 | 0.228 | 0.484 | Del. |  |  |
| 7937335 | IFITM2 | 11 | 2.179 | 0.229 | 0.228 | 0.484 | Del. |  |  |
| 7934731 | C1DP...fg | 10 | 0.217 | -0.9 | -1.12 | -0.6 |  |  |  |
| 7934731 | C1DP2 | 10 | 0.217 | -0.9 | -1.12 | -0.6 |  |  |  |
| 7934731 | C1DP3 | 10 | 0.217 | -0.9 | -1.12 | -0.6 |  |  |  |
| 7934731 | C1DP1 | 10 | 0.217 | -0.9 | -1.12 | -0.6 |  |  |  |
| 7934731 | C1DP4 | 10 | 0.217 | -0.9 | -1.12 | -0.6 |  |  |  |
| 7934731 | C1D | 2 | 0.217 | -0.9 | -1.12 | -0.6 |  |  |  |
| 7903717 | MIR197 | 1 | 0.687 | 1.372 | 1.049 | 0.996 |  |  |  |
| 7952205 | MCAM | 11 | 0.958 | 0.824 | 0.882 | 0.886 | Del. |  |  |
| 7894185 | OAZ1 | 19 | -0.71 | -1.08 | -0.69 | -0.81 |  |  |  |
| 8142763 | Nd | 7 | -0.73 | -0.58 | 0.019 | -0.2 | Del. |  |  |
| 7947230 | BDNF | 11 | -1.14 | -1.57 | -1.3 | -1.32 | Del. | Del. | Del. |
| 8135594 | CAV1 | 7 | -1.17 | -1.22 | -1.38 | -1.26 |  |  |  |
| 7902265 | Nd | 1 | 0.946 | 1.285 | 1.087 | 1.098 |  |  |  |
| 7901175 | TSPAN1 | 1 | 1.563 | 1.468 | 1.121 | 1.37 | Del. |  |  |
| 7916493 | PPAP2B | 1 | 0.755 | 0.616 | 0.514 | 0.621 | Amp. |  |  |
| 7894891 | Nd | 2 | 1.25 | 2.188 | 1.987 | 1.758 |  |  |  |
| 7893711 | ABCF1 | 6 | 1.828 | 1.907 | 1.65 | 1.792 |  |  |  |
| 7995320 | Nd | 16 | 1.188 | 1.597 | 1.266 | 1.339 | Amp. |  |  |
| 7995320 | Nd | 16 | 1.188 | 1.597 | 1.266 | 1.339 | Amp. |  |  |
| 7995320 | Nd | 16 | 1.188 | 1.597 | 1.266 | 1.339 | Amp. |  |  |
| 7995320 | Nd | 16 | 1.188 | 1.597 | 1.266 | 1.339 | Amp. |  |  |
| 7895508 | Nd | 6 | 0.357 | 0.815 | 0.685 | 0.584 |  |  |  |
| 8155497 | FAM27C | 9 | 1.575 | 1.948 | 1.795 | 1.766 | Amp. |  |  |
| 7921987 | TMCO1 | 1 | -0.6 | -0.88 | -0.61 | -0.69 | Del. |  |  |
| 8083453 | Nd | 17 | 0.612 | 0.832 | 0.776 | 0.734 |  |  |  |
| 8083453 | Nd | 17 | 0.612 | 0.832 | 0.776 | 0.734 |  |  |  |
| 8083453 | Nd | 17 | 0.612 | 0.832 | 0.776 | 0.734 |  |  |  |
| 8083453 | Nd | 17 | 0.612 | 0.832 | 0.776 | 0.734 |  |  |  |
| 8083453 | .nd | 2 | 0.612 | 0.832 | 0.776 | 0.734 |  |  |  |
| 8083453 | Nd | 2 | 0.612 | 0.832 | 0.776 | 0.734 |  |  |  |
| 8083453 | Nd | 2 | 0.612 | 0.832 | 0.776 | 0.734 |  |  |  |
| 8083453 | Nd | 2 | 0.612 | 0.832 | 0.776 | 0.734 |  |  |  |
| 8083453 | Nd | 3 | 0.612 | 0.832 | 0.776 | 0.734 |  |  |  |
| 8083453 | Nd | 3 | 0.612 | 0.832 | 0.776 | 0.734 |  |  |  |
| 8083453 | Nd | 3 | 0.612 | 0.832 | 0.776 | 0.734 |  |  |  |
| 8111255 | CDH10 | 5 | 0.53 | 0.763 | 0.896 | 0.713 |  | Amp. |  |
| 7896217 | Nd | 19 | -0.35 | -1.17 | -0.48 | -0.58 |  |  |  |
| 8132962 | CCT6A | 7 | -0.04 | -2.01 | -0.52 | -0.35 | Del. |  |  |
| 8132962 | SNORA15 | 7 | -0.04 | -2.01 | -0.52 | -0.35 | Del. |  |  |
| 7893844 | Nd | 14 | 0.813 | 1.207 | 0.819 | 0.93 |  |  |  |
| 8044080 | SLC9A2 | 2 | -0.85 | -0.7 | -0.73 | -0.76 | Amp. |  |  |
| 8130499 | DYNLT1 | 6 | -0.83 | -1.05 | -1.02 | -0.96 | Del. | Del. |  |
| 8065082 | Nd | 20 | -0.54 | 0.106 | -0.26 | -0.25 |  |  |  |
| 8106923 | NR2F1 | 5 | -0.87 | -0.73 | -0.89 | -0.83 | Del. |  |  |
| 8097256 | FGF2 | 4 | 0.977 | 1.204 | 1.078 | 1.083 |  |  |  |
| 8144667 | SUB1P1 | 8 | -0.68 | -1.04 | -0.79 | -0.83 | Del. |  |  |
| 8082607 | ATP2C1 | 3 | -0.86 | -0.97 | -0.85 | -0.89 | Del. |  |  |
| 7895711 | Nd | 2 | 1.345 | -0.05 | 0.307 | 0.282 |  |  |  |
| 7912994 | IFFO2 | 1 | 1.219 | 0.709 | 0.66 | 0.829 | Del. |  |  |
| 7925531 | AKT3 | 1 | 1.595 | 1.035 | 1.077 | 1.212 | Amp. | Del. |  |
| 7893864 | Nd | 6 | 0.227 | -0.68 | -0.55 | -0.44 |  |  |  |
| 7971669 | Nd | 13 | 0.7 | 1.23 | 0.983 | 0.946 | Del. | Del. | Del. |
| 7895521 | HNRNPD | 4 | -0.61 | -0.74 | -0.29 | -0.51 |  |  |  |
| 7896540 | Nd | 12 | 1.524 | 1.961 | 1.978 | 1.808 |  |  |  |
| 8079426 | TMIE | 3 | 0.318 | 0.756 | 0.443 | 0.474 | Del. |  |  |
| 7895791 | Nd | 19 | -0.69 | -1.01 | -0.15 | -0.47 |  |  |  |
| 7896112 | Nd | 2 | -0.55 | -1.16 | -0.31 | -0.58 |  |  |  |
| 7896112 | IK | 5 | -0.55 | -1.16 | -0.31 | -0.58 |  |  |  |
| 7892996 | Nd | 2 | 0.13 | -0.82 | -0.44 | -0.36 |  |  |  |
| 7892996 | Nd | 5 | 0.13 | -0.82 | -0.44 | -0.36 |  |  |  |
| 8114396 | CDC23 | 5 | -0.69 | -1.1 | -0.67 | -0.8 | Del. |  |  |
| 8100376 | Nd | 4 | 0.755 | 0.991 | 0.717 | 0.813 | Amp. |  |  |
| 7893051 | Nd | 5 | 1.731 | 2.423 | 2.256 | 2.115 |  |  |  |
| 8109424 | Nd | 5 | 1.109 | 1.602 | 1.549 | 1.402 |  |  |  |
| 8105612 | CWC27 | 5 | -0.66 | -0.92 | -0.73 | -0.76 | Amp. |  |  |
| 7905444 | SNX27 | 1 | -0.49 | -0.68 | -0.52 | -0.56 |  |  |  |
| 8052370 | Nd | 2 | 0.843 | 1.339 | 0.915 | 1.011 | Amp. |  |  |
| 8098246 | ANXA10 | 4 | -1.49 | -1.67 | -1.5 | -1.55 | Amp. |  |  |
| 7895085 | SMNDC1 | 10 | 0.287 | -0.72 | -0.84 | -0.56 |  |  |  |

**Supplemental Table S5D**

| **Gene ID** | **Gen symbol** | **Chr.** | **R4.1 logFC** | **R4.2 logFC** | **R4.3 logFC** | **logFC Mean** | **R4.1**  **Copy No.** | **R4.2 Copy No.** | **R4.3**  **Copy No.** |
| --- | --- | --- | --- | --- | --- | --- | --- | --- | --- |
| 8148040 | MAL2 | 8 | -5.55 | -5.56 | -5.68 | -5.6 |  |  |  |
| 8067140 | CYP24A1 | 20 | -5.5 | -5.61 | -6.22 | -5.77 |  |  |  |
| 8148280 | SQLE | 8 | -2.41 | -2.77 | -2.47 | -2.55 |  |  |  |
| 8030804 | CD33 | 19 | 1.24 | 1.81 | 1.768 | 1.586 | Amp. |  | Amp. |
| 7983650 | SLC27A2 | 15 | -3.43 | -3.35 | -2.95 | -3.24 |  |  |  |
| 7960143 | ZNF84 | 12 | 0.19 | -1.85 | -0.5 | -0.56 |  |  |  |
| 8113512 | EPB41L4A | 5 | 2.47 | 2.06 | 2.797 | 2.421 |  | Amp. |  |
| 8055496 | LRP1B | 2 | 2.02 | 0.89 | 2.048 | 1.544 | Amp. | Amp. | Amp. |
| 8135763 | WNT16 | 7 | -0.33 | -1.45 | -1.41 | -0.88 |  |  |  |
| 8129476 | C6orf191 | 6 | 0.67 | 0.83 | 2.264 | 1.076 |  |  |  |
| 8098246 | ANXA10 | 4 | -1.82 | -1.3 | -1.2 | -1.42 |  |  |  |
| 7916862 | WLS | 1 | 0.91 | 0.94 | 1.253 | 1.025 |  |  |  |
| 8135587 | CAV2 | 7 | -1.53 | -1.2 | -1.53 | -1.41 |  |  |  |
| 8172158 | CASK | X | -2.04 | -2.02 | -1.96 | -2.01 |  |  | Del. |
| 8023561 | LMAN1 | 18 | -3.1 | -3.36 | -3.05 | -3.17 |  | Amp. | Amp. |
| 7901175 | TSPAN1 | 1 | 0.72 | 1.65 | 0.988 | 1.054 |  |  |  |
| 8036318 | ZNF566 | 19 | 1.19 | -0.44 | 1.368 | 0.893 |  |  |  |
| 7961166 | KLRC4 | 12 | 0.38 | -0.72 | 1.128 | 0.677 |  |  |  |
| 8115327 | SPARC | 5 | 2.8 | 2.76 | 2.87 | 2.809 |  |  |  |
| 8148309 | ND | 8 | -1.33 | -2 | -1.34 | -1.53 |  |  |  |
| 8103415 | FAM198B | 4 | 0.96 | 1.29 | 2.959 | 1.544 |  |  |  |
| 8028058 | KIRREL2 | 19 | 1.54 | 1.43 | 1.52 | 1.494 |  |  |  |
| 8135594 | CAV1 | 7 | -2.22 | -1.89 | -2.22 | -2.1 |  |  |  |
| 8151496 | ZNF704 | 8 | 1.4 | 1.03 | 1.118 | 1.174 |  |  |  |
| 8102415 | CAMK2D | 4 | -1.59 | -1.38 | -1.54 | -1.5 | Del. |  |  |
| 8038192 | FUT1 | 19 | 0.58 | 1.2 | 0.358 | 0.629 |  |  |  |
| 8166747 | SYTL5 | X | -1.53 | -1.63 | -2.13 | -1.74 |  |  |  |
| 8106986 | RHOBTB3 | 5 | -0.86 | -1.59 | -0.8 | -1.03 |  |  |  |
| 7977933 | SLC7A8 | 14 | 1.27 | 1.11 | 1.885 | 1.385 | Amp. |  | Amp. |
| 7902104 | PDE4B | 1 | -1.56 | -1.81 | -1.36 | -1.57 |  |  |  |
| 8003060 | SDR42E1 | 16 | -1.4 | -1.46 | -1.2 | -1.35 |  |  |  |
| 7954559 | PPFIBP1 | 12 | 0.14 | -1.05 | 0.143 | -0.28 |  |  |  |
| 8138805 | CPVL | 7 | 1.11 | 0.64 | 0.932 | 0.872 |  |  |  |
| 8180200 | ZNF493 | 19 | -0.77 | -0.72 | -1.11 | -0.85 |  |  |  |
| 7934970 | HTR7 | 10 | -1.28 | -1.21 | -1.59 | -1.35 |  |  |  |
| 7932744 | ARMC4 | 10 | 0.23 | -0.9 | 0.348 | -0.42 |  |  |  |
| 8072587 | SLC5A1 | 22 | 0.34 | 0.75 | 1.506 | 0.73 |  |  |  |
| 8096160 | ARHGAP24 | 4 | 1.26 | 1.28 | 1.282 | 1.276 | Del. |  |  |
| 7982066 | Nd | 15 | -0.12 | 2.09 | 0.734 | 0.568 | Amp. |  | Amp. |
| 7982066 | SNORD115-24 | 15 | -0.12 | 2.09 | 0.734 | 0.568 | Amp. |  | Amp. |
| 7982066 | SNORD115-30 | 15 | -0.12 | 2.09 | 0.734 | 0.568 | Amp. |  | Amp. |
| 7982066 | SNORD115-42 | 15 | -0.12 | 2.09 | 0.734 | 0.568 | Amp. |  | Amp. |
| 7978376 | STXBP6 | 14 | -0.66 | 0.06 | -0.88 | -0.33 | Amp. | Amp. | Amp. |
| 8127563 | COL12A1 | 6 | -0.83 | -1.61 | -1.24 | -1.18 |  | Amp. |  |
| 8035847 | ZNF675 | 19 | -0.62 | -1.4 | -0.5 | -0.76 | Amp. |  | Amp. |
| 8069880 | TIAM1 | 21 | -0.88 | -0.8 | -1.03 | -0.9 |  |  |  |
| 8126820 | GPR110 | 6 | -0.4 | -1.56 | 0.481 | -0.67 |  |  |  |
| 8040163 | IAH1 | 2 | -0.86 | -0.89 | -0.99 | -0.91 |  |  |  |
| 8099393 | Nd | 4 | -1.23 | -0.22 | -0.75 | -0.58 |  | Amp. |  |
| 7926875 | BAMBI | 10 | 0.42 | 1.32 | 1.625 | 0.964 |  |  |  |
| 8081214 | GPR15 | 3 | -1.24 | -1.54 | -1.3 | -1.36 |  |  |  |
| 8167973 | HEPH | X | 1.31 | 0.76 | 0.814 | 0.933 |  |  |  |
| 8110084 | MSX2 | 5 | -1.49 | -1.35 | -1.44 | -1.43 |  |  |  |
| 8174527 | CAPN6 | X | 0.96 | 0.68 | 1.222 | 0.929 |  |  |  |
| 7943263 | AMOTL1 | 11 | 0.29 | -0.79 | -0.05 | -0.23 |  |  |  |
| 8149927 | CLU | 8 | -0.43 | -0.66 | -0.73 | -0.59 |  |  |  |
| 8085263 | TMEM111 | 3 | -1.23 | -1.27 | -1.3 | -1.26 |  |  |  |
| 7960134 | ZNF26 | 12 | -1.58 | -1.82 | -1.32 | -1.56 |  |  |  |
| 8175217 | GPC4 | X | -0.5 | 0.77 | 0.551 | 0.595 |  |  |  |
| 7951077 | SESN3 | 11 | -1.87 | -1.9 | -1.31 | -1.67 |  |  |  |
| 8117045 | RBM24 | 6 | 0.32 | -1.09 | -0.22 | -0.43 | Amp. |  | Amp. |
| 8053325 | Nd | 2 | 0.34 | 0.99 | 1.27 | 0.754 |  |  |  |
| 7961175 | KLRC3 | 12 | -0.09 | -0.79 | 0.38 | -0.3 |  |  |  |
| 8168749 | SRPX2 | X | -0.93 | -0.89 | -1.23 | -1 |  |  |  |
| 7932765 | MPP7 | 10 | 0.07 | -1.14 | -0.2 | -0.26 | Del. | Del. | Del. |
| 8060988 | BTBD3 | 20 | 1.37 | 1.16 | 1.154 | 1.222 |  |  |  |
| 8049487 | MLPH | 2 | -1.17 | -1.22 | -1.38 | -1.25 | Amp. | Amp. | Amp. |
| 8035842 | ZNF91 | 19 | -0.41 | -1.51 | -1.06 | -0.87 |  |  | Amp. |
| 8033754 | ZNF266 | 19 | -1.4 | -1.19 | -1.22 | -1.27 |  |  |  |
| 8062041 | ACSS2 | 20 | 0.52 | 1.22 | 0.291 | 0.568 |  |  |  |
| 7997010 | CLEC18...fg | 16 | -0.95 | 0.29 | -1.55 | -0.75 |  |  | Amp. |
| 7997010 | CLEC18A | 16 | -0.95 | 0.29 | -1.55 | -0.75 |  |  | Amp. |
| 7997010 | CLEC18C | 16 | -0.95 | 0.29 | -1.55 | -0.75 |  |  | Amp. |
| 8015133 | KRT23 | 17 | -2.08 | -1.84 | -0.81 | -1.46 | Amp. |  | Amp. |
| 8074853 | ZNF280A | 22 | -0.78 | -0.65 | -0.77 | -0.73 |  |  |  |
| 7958352 | BTBD11 | 12 | 1.19 | 1.37 | 1.502 | 1.349 |  |  |  |
| 7951686 | IL18 | 11 | -0.85 | 0.11 | -0.08 | -0.19 |  |  |  |
| 8175269 | FAM122B | X | -0.7 | -0.6 | -0.55 | -0.61 |  |  |  |
| 8045336 | GPR39 | 2 | 0.29 | 1.34 | -0.07 | 0.301 | Del. | Del. | Del. |
| 7960529 | SCNN1A | 12 | -0.98 | -0.23 | -1.11 | -0.63 |  |  |  |
| 7896179 | Nd | 14 | -0.16 | -1.04 | 0.045 | -0.2 |  |  |  |
| 8161737 | Nd | 9 | -0.74 | -1.09 | -0.64 | -0.8 | Del. | Del. | Del. |
| 8117415 | HIST1H3E | 6 | 0.65 | 0.56 | 0.808 | 0.665 | Amp. |  | Amp. |
| 8145365 | DOCK5 | 8 | -0.89 | -0.46 | -0.73 | -0.67 |  |  |  |
| 8063923 | SLCO4A1 | 20 | 1.07 | 1.14 | 0.805 | 0.995 | Amp. |  |  |
| 7961151 | KLRK1 | 12 | 0.42 | -0.32 | 1.368 | 0.567 |  |  |  |
| 7893748 | Nd | 16 | -0.42 | -0 | 0.633 | 0.096 |  |  |  |
| 8150862 | Nd | 8 | -0.78 | -0.85 | -0.86 | -0.83 |  |  |  |
| 7951036 | SNORD5 | 11 | -0.86 | -1.07 | -0.83 | -0.91 |  |  |  |
| 7951036 | SNORA18 | 11 | -0.86 | -1.07 | -0.83 | -0.91 |  |  |  |
| 7951036 | MIR1304 | 11 | -0.86 | -1.07 | -0.83 | -0.91 |  |  |  |
| 8082058 | CSTA | 3 | -0.01 | 1.55 | -0.06 | 0.083 |  |  |  |
| 7966690 | TBX3 | 12 | 1.25 | 0.36 | 1.135 | 0.802 | Del. | Del. | Del. |
| 7894895 | ILF2 | 1 | -1.42 | -0.49 | 0.484 | -0.7 |  |  |  |
| 8035318 | UNC13A | 19 | 0.46 | 0.83 | 0.616 | 0.618 | Amp. |  | Amp. |
| 8134219 | CCDC132 | 7 | -0.83 | -0.76 | -0.5 | -0.68 |  |  |  |
| 8106727 | ATP6AP1L | 5 | -0 | 1.25 | 0.322 | 0.12 |  |  |  |
| 8140668 | SEMA3A | 7 | 0.83 | 0.53 | 1.002 | 0.762 |  |  |  |
| 8103563 | DDX60 | 4 | -0.58 | -0.34 | 0.693 | -0.52 |  |  |  |
| 8098441 | ODZ3 | 4 | -0.86 | -0.9 | -0.73 | -0.82 |  |  |  |

**Supplemental Table S6**

| **Gene** | **p53+/+: CYC116**  **clones** | | **p53-/-:CYC116**  **clones** | | **p53+/+: ZM447439**  **clones** | | **p53-/-: ZM447439**  **clones** | |
| --- | --- | --- | --- | --- | --- | --- | --- | --- |
|  | Microarray | qRT-PCR | Microarray | qRT-PCR | Microarray | qRT-PCR | Microarray | qRT-PCR |
| CYP24A1 | -32 | -33 | -30 | -50 | NC^a^ | NC | -55 | -200 |
| Bcl-xL | 2 | 2 | 1.5 | 2 | NC | NC | NC | NC |
| GJC1 | -3 | -3.5 | -5 | -5 | NC | NC | NC | NC |
| NCAM1 | 2 | 22 | NC | NC | NC | NC | NC | NC |
| KLK5 | 2.34 | 62 | NC | NC | NC | NC | NC | NC |
| KRT7 | 2 | 30 | NC | NC | NC | NC | NC | NC |
| LCN2 | 7 | 229 | NC | NC | NC | NC | NC | NC |
| TNFAIP3 | 3.22 | 11 | NC | NC | NC | NC | NC | NC |
| KRT13 | 3.4 | 396 | NC | NC | NC | NC | NC | NC |
| PPAP2B | 1.4 | 7 | NC | NC | 2 | 2.3 | NC | NC |
| TBX3 | 3.3 | 11 | NC | NC | NC | NC | 2 | 7.4 |
| SERINC2 | 5 | 7.4 | NC | NC | 2 | 2.1 | NC | NC |
| HOXB5 | 4 | 5.4 | NC | NC | NC | NC | NC | NC |
| ANXA10 | -2 | -2 | NC | NC | -3 | -6 | -3 | -1.3 |
| CYP1A1 | 3 | 9 | 8 | 28 | NC | NC | NC | NC |
| PRKACB | -9 | -6 | -3 | -3 | NC | NC | NC | NC |
| A4GALT | 3 | 6.3 | 2.3 | 6.2 | NC | NC | NC | NC |
| ARHGAP29 | -5 | -5 | -4.3 | -2.3 | NC | NC | NC | NC |
| NRP1 | 1.4 | 14 | NC | NC | NC | NC | NC | NC |
| KLRK1 | -4 | -3 | NC | NC | NC | NC | 1.5 | 1.3 |
| MID1 | NC | NC | -18 | -3 | NC | NC | NC | NC |
| EHF | NC | NC | 8.4 | 264 | NC | NC | NC | NC |
| SEMA3A | NC | NC | **-2** | **3** | NC | NC | 2 | 3 |
| PLK4 | NC | NC | -2.2 | -1.1 | NC | NC | NC | NC |
| INPP4B | NC | NC | -2.4 | -1.2 | NC | NC | NC | NC |
| CAMK2D | NC | NC | -2 | -1.4 | NC | NC | -3 | -3.2 |
| BDNF | NC | NC | -1.5 | -1.4 | -2.5 | -4.5 | NC | NC |
| TSPAN1 | NC | NC | NC | NC | 2.6 | 2 | 2.1 | 4 |

**Supplemental Table S7**

| **Group** | **Common pathways** | **Differential pathways** |
| --- | --- | --- |
| p53+/+ and p53-/-  CYC116 resistant clones | Cell adhesion-alpha‑4 integrins  in cell migration and adhesion,  signal transduction-Erk  Interactions, signal transduction-cAMP signaling, transport-ACM3  in salivary glands, and regulation  of lipid metabolism-regulation of  lipid metabolism by niacin and  Isoprenaline. | DNA damage-mismatch repair,  cell cycle-Spindle assembly and  chromosome separation, and cell  cycle-role of APC in cell cycle  regulation. |
| p53+/+ and p53-/-  ZM447439 resistant clones | Delta508‑CFTR traffic/  ER‑to‑golgi, normal wtCFTR  traffic/ER‑to‑golgi,  neurophysiological process-  NMDA‑dependent postsynaptic  long‑term potentiation in CA1  hippocampal neurons,  neurophysiological process-  dopamine D2 receptor  transactivation of PDGFR in  CNS, and cholesterol and  sphingolipids transport/Influx  to the early endosome in lung. | Immune response-classical  complement pathway and immune  response-human NKG2D signaling. |

**Supplemental materials and methods**

**Computational modeling**

**ZM447439 in complex with Aurora B kinase**

The X-ray crystal structure of Aurora B kinase in complex with ZM447439 was taken from the PDB database (code 2VRX) (1) The complex was prepared in the following way: hydrogens were added to the protein using the Reduce (2) and LEaP programs (3) and to the ligands using Chimera, ver. 1.5.3 (4). The parameters for the protein were acquired from the ff03 force field (5) and for the ligands from the gaff force field (6). Charges for the ligands were calculated using the RESP procedure (7) at the HF/6-31G* level. The complex was relaxed in several steps. First, the hydrogens were optimized using AMBER program (3) for 2000 steps followed by a short high-temperature molecular dynamics (1 ps, starting from 1700 K, cooled down to 10 K).

Based on this structure, three different mutant proteins (mutant-1: I216L; mutant-2: N76V, L152S; mutant-3: L152S) were built automatically using the LEaP (3) program and adjusted manually by use of Pymol (8). Subsequently, the modeled residues were relaxed using AMBER minimization for 5000 steps, followed by 1 ps molecular dynamics (with three independent calculations starting at 300, 600 or 1200 K; and cooled down to 10 K). However, all molecular dynamics run resulted into almost identical geometries and hence only one of them was used for the optimization. The relaxed mutant proteins were further treated as the wild-type protein.

The protein-inhibitor interaction energies were calculated using our SQM/MM procedure (semiempirical quantum chemistry linked to molecular mechanics) for optimization and scoring (9,10). The SQM part comprised the ligand and the amino acids of the protein extending to 6 Å from the ligand. The rest of the protein was calculated using MM (AMBER) and kept frozen. The surrounding was modeled using the generalized Born (GB) solvation model mimicking the solvent (11). All complexes were optimized in a SQM/MM setup using our in-house program linking the SQM program (MOPAC2009) and the MM program (AMBER) (3). The SQM part was treated by the newly parametrized PM6-D3H4X method (12) which was shown to reproduce experimental binding constants closely (10). The MOZYME approximation was used to speed up the calculations. The SQM/MM optimizations were performed in several rounds until the energy and gradient convergence criteria (ΔE = 0.005 kcal/mol, maximum gradient of 1 kcal/mol/Å, root-mean-square of the gradient of 0.5 kcal/mol/Å) were met. The SQM/MM optimized structures were subsequently scored using our recently developed scoring methods (9).

**Docking of CYC116 in Aurora B kinase using Glide and SQM/MM rescoring**

Aurora B kinase complex (code 2VRX) was subjected to preparation steps using the Protein Preparation Wizard in Maestro (13): waters were removed, bond orders were assigned and hydrogens were added. Next, the orientation of amide (Asn and Gln), hydroxyl (Ser, Thr, and Tyr), and thiol groups (Cys) and the protonation and tautomeric state of His residues were optimized using the exhaustive sampling option. For generation of receptor grids, a grid box of 20 × 20 × 20 Å3 with a default inner box (10 × 10 × 10 Å3) was centered on the corresponding ligand. Default parameters were used, and no constraints were included. Docking calculations were performed using Glide Extra Precision (XP) (14) algorithm along with postdocking minimization introduced as default in the Glide 5.5 for XP docking as postprocessing. In the protocol, Glide was set to write out the 5 best poses per ligand. The best pose was reoptimized and rescored using our SQM/MM-based (PM6-D3H4X) scoring function in the manner analogous to that in case of ZM447439.

**pH3Ser10 staining**

The cells were harvested and fixed following the cell cycle method as described in the main text. The cells were washed in PBS, 1% fetal bovine serum (FBS) solution. The pellet was suspended in PBS, 0.25% Triton X-100 (Sigma) and incubated on ice for 15 min. The pellet was washed with PBS, 1% FBS solution and stained with 100 µl of phospho-histoneH3 antibody (Upstate 1:500) for 1 h. The unbound antibody was washed out and suspended in 100 µl of secondary Alexa flour 488 goat anti-rabbit IgG (Invitrogen, 1:500) and incubated for 30 min. After washing, the pellet was suspended in DNA staining/RNase solution (50 µg/ml propidium iodide, 0.5 mg/ml RNase in PBS, 1%FBS) and incubated in dark at 37°C for 30 min and finally analyzed by flowcytometry (FACSCalibur, Becton Dickinson).

**Cytogenetic arrays**

DNA was extracted from one million cells using DNeasy blood and tissue kit (QIAGEN). Extracted genomic DNA was processed exactly according to manufacturer’s protocol (Affymetrix, Santa Clara, CA). 100 ng of DNA was amplified by whole genome amplification. After product purification with magnetic beads, DNA was quantified, fragmented, labeled, and hybridized to Cytogenetics Whole-Genome 2.7M array. Arrays were washed, stained and scanned. We used software Partek Genomics Suite to analyze CGH arrays. Corresponding copy number changes for differentially expressed genes (p<0.001) were shown in the circos plots and also in the supplemetary tables S5, S6, S7, and S8.
